# Cryo-EM structure of the complete inner kinetochore of the budding yeast point centromere

**DOI:** 10.1101/2022.12.12.520091

**Authors:** Tom Dendooven, Ziguo Zhang, Jing Yang, Stephen H. McLaughlin, Johannes Schwab, Sjors H.W. Scheres, Stanislau Yatskevich, David Barford

**Author notes:** These authors contributed equally (Tom Dendooven, Ziguo Zhang).

## Abstract

**Summary:** The point centromere of budding yeast specifies assembly of the large multi-subunit kinetochore complex. By direct attachment to the mitotic spindle, kinetochores couple the forces of microtubule dynamics to power chromatid segregation at mitosis. Kinetochores share a conserved architecture comprising the centromere-associated inner kinetochore CCAN (constitutive centromere-associated network) complex and the microtubule-binding outer kinetochore KMN network. The budding yeast inner kinetochore additionally includes the centromere-binding CBF1 and CBF3 complexes. Here, we reconstituted the complete yeast inner kinetochore complex assembled onto the centromere-specific CENP-A nucleosome (CENP-A^Nuc^) and determined its structure using cryo-EM. This revealed a central CENP-A^Nuc^, wrapped by only one turn of DNA, and harboring extensively unwrapped DNA ends. These free DNA duplexes function as binding sites for two CCAN protomers, one of which entraps DNA topologically and is positioned precisely on the centromere by the sequence-specific DNA-binding complex CBF1. The CCAN protomers are connected through CBF3 to form an arch-like configuration, binding 150 bp of DNA. We also define a structural model for a CENP-A^Nuc^-pathway to the outer kinetochore involving only CENP-QU. This study presents a framework for understanding the basis of complete inner kinetochore assembly onto a point centromere, and how it organizes the outer kinetochore for robust chromosome attachment to the mitotic spindle.

## Introduction

The survival of all living species depends on the faithful inheritance of their genetic information. Eukaryotes achieve this using a microtubule-based spindle apparatus to segregate replicated sister chromatids at mitosis. Critical to this process is the kinetochore, a large multi-subunit complex that attaches chromatids to the mitotic spindle, and harnesses the power of microtubule depolymerization to move chromatids to opposite ends of the spindle (Musacchio and Desai, 2017; Navarro and Cheeseman, 2021; Sridhar and Fukagawa, 2022). In many organisms, kinetochore assembly is restricted to the centromere, a specialized region of chromatin defined by nucleosomes containing the histone H3 variant CENP-A (CENP-A^Nuc^) (Earnshaw and Rothfield, 1985; Meluh et al., 1998; Stoler et al., 1995). Kinetochores are structurally and functionally delineated into the inner and outer kinetochore. The inner kinetochore CCAN complex, also referred to as the Ctf19 complex in *S. cerevisiae* (Cheeseman et al., 2002), associates with the centromere, generally through specific recognition of CENP-A^Nuc^. CCAN then connects the centromere to the outer kinetochore - the ten-subunit KMN network. Ndc80 of this network, in association with either the Dam1/DASH complex in yeast, or the Ska complex in humans, attaches kinetochores to spindle microtubules.

The mechanisms of selective CCAN stoichiometry and assembly onto centromeric chromatin, and how it forms connections to the outer kinetochore, are long-standing questions. CCAN is a 13-16 subunit complex composed of distinct sub-modules (**Table S1**). Most CCAN genes are essential in humans, whereas in budding yeast only CENP-C, CENP-Q and CENP-U are absolutely required for viability (Brown et al., 1993; Cheeseman et al., 2002; Meluh and Koshland, 1995). Disruption of most other CCAN genes introduces chromosome segregation defects, and all CCAN genes are required for budding yeast meiosis (Borek et al., 2021). In animals, the interactions of CENP-C and CENP-N with CENP-A^Nuc^ determines the selectivity of CCAN for centromeres (Carroll et al., 2010; Carroll et al., 2009). *S. cerevisiae* CENP-C also binds CENP-A^Nuc^, and is required for centromere recruitment of some CCAN subunits, (Cohen et al., 2008; Hornung et al., 2014; Meluh and Koshland, 1995; Westermann et al., 2003; Xiao et al., 2017). Specific to budding yeast, CENP-QU associates with an essential N-terminal domain of CENP-A (CENP-A^END^) (Anedchenko et al., 2019; Chen et al., 2000; Fischbock-Halwachs et al., 2019; Keith et al., 1999).

The point centromeres of budding yeast and regional centromeres of higher eukaryotes differ substantially in size and higher-order structure but nevertheless share a conserved underlying architecture. At a structural level, point centromeres comprise an individual CENP-A^Nuc^-kinetochore complex that attaches to a single microtubule (Furuyama and Biggins, 2007; Winey et al., 1995). The budding yeast centromere is genetically defined by a ∼120 bp sequence that is sufficient to template complete mitotic and meiotic centromere function (Cottarel et al., 1989), and onto which CENP-A^Nuc^ is perfectly positioned (Cole et al., 2011; Krassovsky et al., 2012). All 16 *S. cerevisiae* centromeres comprise three centromere DNA elements (CDE) (**Figure 1A**) (Clarke and Carbon, 1980, 1983; Fitzgerald-Hayes et al., 1982). The two short CDEI and CDEIII motifs are highly conserved, and function to bind CBF1 and CBF3 respectively, two protein complexes that are specific to the point centromere-kinetochores of budding yeast (Baker and Masison, 1990; Cai and Davis, 1990; Lechner and Carbon, 1991; Mellor et al., 1990). Whereas CDEIII and CBF3 are essential for viability (Doheny et al., 1993; Goh and Kilmartin, 1993; Hegemann et al., 1988; Jehn et al., 1991; Lechner and Carbon, 1991; McGrew et al., 1986; Ng and Carbon, 1987; Panzeri et al., 1985), cells with CDEI disrupted remain viable but exhibit mitotic chromosome loss, and defective centromere function in meiosis I (Cumberledge and Carbon, 1987; Gaudet and Fitzgerald-Hayes, 1989; Hegemann et al., 1988; Niedenthal et al., 1991; Panzeri et al., 1985). CDEII is less well conserved, however, its AT-rich DNA sequence is proposed to be favorable for CENP-A^Nuc^ wrapping due to its increased tendency to curve (Bechert et al., 1999; Koo et al., 1986; Murphy et al., 1991; Ortiz et al., 1999). Indeed, CDEII is necessary for optimal centromere function: reduction of AT-content, disruption of polyA/T tracts and alteration to CDEII length, result in mitotic delay and chromosome segregation defects (Baker and Rogers, 2005; Cumberledge and Carbon, 1987; Gaudet and Fitzgerald-Hayes, 1987, 1989; Murphy et al., 1991; Spencer and Hieter, 1992).

**Figure 1.**
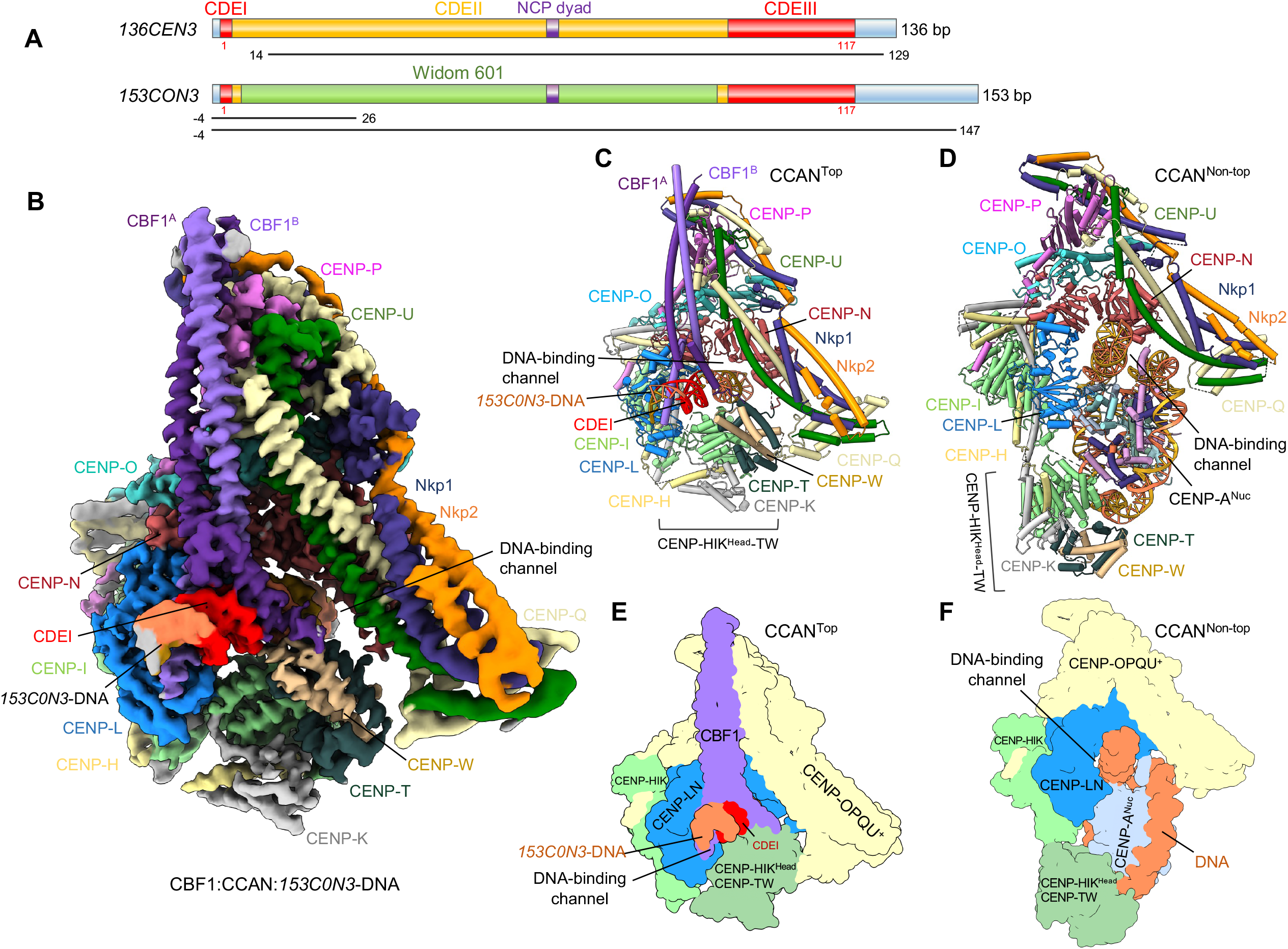
CBF1 is a bHLH homo-dimer that positions the CBF1:CCAN complex onto CDEI. (**A**) Schematic of the native *136CEN3* and chimeric *153C0N3* sequences discussed in this study. In the *153C0N3* sequence, a large part of CDEII is replaced by the Widom 601 sequence to increase nucleosome stability. The regions of *136CEN3* that interact with the CENP-A histone octamer in the native *CEN3*-CENP-A^Nuc^ structure (Guan et al., 2021), and *153C0N3* that interacts with CBF1:CCAN and the inner kinetochore (this study) are indicated below as black lines. (**B**) Cryo-EM map of the CBF1:CCAN complex bound to a 30 bp DNA segment of *153C0N3* containing CDEI, at 3.4 Å. **(C)** Ribbons representation of the CBF1:CCAN:DNA complex with components annotated and a schematic below (**E**). The CENP-HIK^Head^-TW module topologically entraps the *CEN3* DNA within the CENP-LN DNA-binding channel. The CENP-HIK^Head^-TW module adopts an ‘upwards’ conformation, contacting the DNA duplex. (**D**) Structure of CCAN:*W601*-CENP-A^Nuc^ complex with CENP-A^Nuc^ reconstituted with Widom 601 DNA, in the absence of CBF1. Schematic below (**F**). From (Yan et al., 2019). CENP-HIK^Head^-TW module adopts an ‘downwards’ conformation, interacting with the CENP-A^Nuc^ DNA gyre.

How CBF1 recognizes CDEI and its role in assembling a complete kinetochore at point centromeres remains unknown. The effects of deleting either CDEI or CBF1 on chromosome segregation fidelity during mitosis are similar (Baker and Masison, 1990; Cai and Davis, 1990; Mellor et al., 1990), indicating that CBF1 is the key mediator of CDEI function. CBF3 initiates *S. cerevisiae* kinetochore assembly by mediating CENP-A^Nuc^ deposition at centromeric loci (Camahort et al., 2007; Cho and Harrison, 2011; Shivaraju et al., 2011). However, since CBF3 remains localized at the centromere during metaphase and anaphase (Cieslinski et al., 2021), it likely has a structural role in kinetochore assembly beyond CENP-A^Nuc^ deposition.

The findings described above paint a molecular picture of the yeast inner kinetochore consisting of CCAN, CBF1 and CBF3 assembled onto CENP-A^Nuc^:CENP-C (Akiyoshi et al., 2009; Cheeseman et al., 2002; De Wulf et al., 2003; Ortiz et al., 1999). We previously determined a cryo-EM structure of *S. cerevisiae* CCAN in complex with CENP-A^Nuc^ reconstituted with non-native Widom 601 DNA (Yan et al., 2019). The structure delineated the overall architecture of CCAN and revealed an interaction between the unwrapped DNA terminus of CENP-A^Nuc^ and a deep positively-charged DNA-binding channel situated at the center of CCAN. Here, we build on previous studies by solving the structure of the entire inner kinetochore, incorporating CCAN, CBF1 and CBF3 assembled on CENP-A^Nuc^ wrapped by near-native centromere DNA, and provide *in vivo* support for our models. Both CBF1 and CBF3 function to organize two CCAN protomers onto a central CENP-A^Nuc^. Dimeric CBF1 binds CDEI with its basic-helix-loop-helix segments to position one of the two CCAN protomers 5’ of CENP-A^Nuc^. We describe an alternative DNA-binding mode for this CCAN where CBF1 assists in the topological entrapment of DNA via the CENP-HIK^Head^-TW module of CCAN, reminiscent of how human CCAN entraps α-satellite linker DNA (Yatskevich et al., 2022). A second CCAN assembles onto the 3’-end of the DNA using a non-topological DNA-binding mode identical to our previous CCAN:CENP-A^Nuc^ structure (Yan et al., 2019; Zhang et al., 2020), generating an asymmetric, dimeric-CCAN inner kinetochore. The two CCAN modules are bridged by CBF3^Core^, now displaced from the CENP-A^Nuc^ face (Guan et al., 2021), to fulfil a stabilizing role at the kinetochore. Together, the inner kinetochore forms an arch-like structure around a central CENP-A^Nuc^, embedding ∼150 bp of centromeric DNA.

Finally, we present a structural explanation for how the CENP-A N-terminus (CENP-A^N^) interacts with CENP-QU (Anedchenko et al., 2019; Chen et al., 2000; Fischbock-Halwachs et al., 2019; Keith et al., 1999). Surprisingly, the CENP-A^N^-binding site on CENP-QU is auto-inhibited in the context of the entire CCAN. Thus, CENP-QU binds CENP-A^N^ independently of CCAN, suggesting a separate CENP-A^Nuc^-CENP-QU connection to the outer kinetochore.

## Results

### *In vitro* reconstituted holo-inner kinetochore complex has a mass of 1.6 MDa

In our previous study, we used the strong positioning Widom 601 DNA sequence (Lowary and Widom, 1997) to generate stable CENP-A^Nuc^ (termed *W601*-CENP-A^Nuc^) for reconstituting CCAN:CENP-A^Nuc^ complexes for cryo-EM (Yan et al., 2019). CENP-A^Nuc^ comprising native centromeric DNA (*CEN*-CENP-A^Nuc^) on the other hand, may have different DNA wrapping properties from that reconstituted with the Widom 601 sequence. In addition, the binding of the inner kinetochore complexes CBF1 and CBF3 to specific sequence elements present in *CEN*-CENP-A^Nuc^ but not *W601*-CENP-A^Nuc^, may influence the organization and stoichiometry of CCAN on CENP-A^Nuc^. Because of the instability of *CEN*-CENP-A^Nuc^ (Dechassa et al., 2011; Guan et al., 2021), difficulties assembling stable CCAN complexes on *CEN*-CENP-A^Nuc^ suitable for cryo-EM had frustrated our earlier efforts to understand the inner kinetochore structure. Recently, a single chain antibody fragment (scFv) specific for the H2A-H2B histone dimer was used to stabilize an *S. cerevisiae* centromeric nucleosome (*CEN3*-CENP-A^Nuc^) for cryo-EM analysis (Guan et al., 2021).

The determined structure, at 3.1 Å resolution, defined the position of the *CEN3* DNA on the histone octamer, and revealed that a 20 bp palindrome in *CEN3* is centered exactly on the dyad axis of the histone octamer **(Figures 1A and S1A**). Using this information, we designed a chimeric 153 bp DNA sequence (*153C0N3*) incorporating the CDEI and CDEIII elements, and their flanking sequences, and substituted Widom 601 sequence for most of CDEII **(Figures 1A and S1A**). We used *153C0N3* to generate a native-like, but more stable CENP-A^Nuc^ (*153C0N3*-CENP-A^Nuc^). We then used *153C0N3*-CENP-A^Nuc^ to reconstitute the holo-inner kinetochore complex with CBF1 and CBF3^Core^ (CBF1:CCAN:*153C0N3*-CENP-A^Nuc^:CBF3^Core^) (**Figures S1B and S1C)**. SEC-MALS analysis showed that the holo-inner kinetochore complex had an overall molecular mass of 1.6 MDa, consistent with a stoichiometry of (CBF1)_2_:(CCAN)_2_:*153C0N3*-CENP-A^Nuc^:CBF3^Core^ (**Figure S1D**).

### CBF1 engages CDEI to position a CCAN protomer at the 5’end of the centromere

In our first attempt to determine a cryo-EM structure of the holo-inner kinetochore we observed cryo-EM particles comprising a CBF1:CCAN complex bound to a 30 bp segment of DNA corresponding to the 5’end of *153C0N3* that includes CDEI (**Figures 1A and S2A**). Despite the presence of the stabilizing Widom 601 sequence within *153C0N3*, CENP-A^Nuc^ had likely denatured at the air-water interface of the cryo-EM grids. Nevertheless, the CBF1:CCAN:DNA complex at 3.4 Å resolution (**Figure S3A and Table S2**) revealed how CBF1 binds to a single CCAN promoter, precisely positioning CCAN onto CDEI of centromeric DNA (**Figures 1B, 1C and 1E)**. The CBF1:CCAN complex contacts centromeric DNA within the DNA-binding CENP-LN channel as observed previously for the CCAN:*W601*-CENP-A^Nuc^ complex (Yan et al., 2019) (**Figure 1D, F)**. Only the ordered basic-helix-loop-helix-leucine zipper domain of CBF1 (residues 218-333) (bHLH) is observed in the cryo-EM map. This DNA-binding bHLH is sufficient for CBF1’s chromosome segregation functions in cells (Mellor et al., 1990). One consequence of CBF1 engaging CCAN is that, compared with our previously described CCAN:*W601*-CENP-A^Nuc^ structure (Yan et al., 2019), CBF1 extends the CENP-LN channel, so a total of 30 bp of DNA interact with CBF1:CCAN (**Figures 2A and 2B**). Additionally, in the CBF1:CCAN:DNA complex, the CENP-LN channel converts into an enclosed basic chamber that completely surrounds the DNA duplex (**Figures 2A and 2B**), a configuration strikingly reminiscent of how human CCAN grips the linker DNA of an α-satellite-CENP-A^Nuc^ (Yatskevich et al., 2022). Formation of the enclosed DNA-binding chamber is mediated by the mobile CENP-HIK^Head^-TW module adopting a raised position, relative to CCAN:*W601*-CENP-A^Nuc^ (Yan et al., 2019), to directly contact the DNA duplex (**Figures 1C, 1E and 2A**). Specifically, a basic surface on CENP-I^Head^ forms extensive contacts with the DNA-phosphate back-bone (**Figure 2B**). This topologically enclosed DNA-binding chamber is stabilized through interactions between an acidic patch on CENP-TW with the CBF1^A^ protomer (**Figures 2A and 2B**). In the previously described CCAN:*W601*-CENP-A^Nuc^ structure (Yan et al., 2019), on the other hand, CENP-HIK^Head^-TW contacts the DNA gyre of CENP-A^Nuc^ at SHL0 (**Figures 1D and 1F**). Thus, this new CBF1:CCAN:DNA structure revealed that yeast CCAN employs two modes of binding to centromeric DNA that differ in the position of the CENP-HIK^Head^-TW module, and constitute topological and non-topological DNA-binding mechanisms, referred to as CCAN^Top^ and CCAN^Non-top^, respectively (**Figures 1C-1F**).

**Figure 2.**
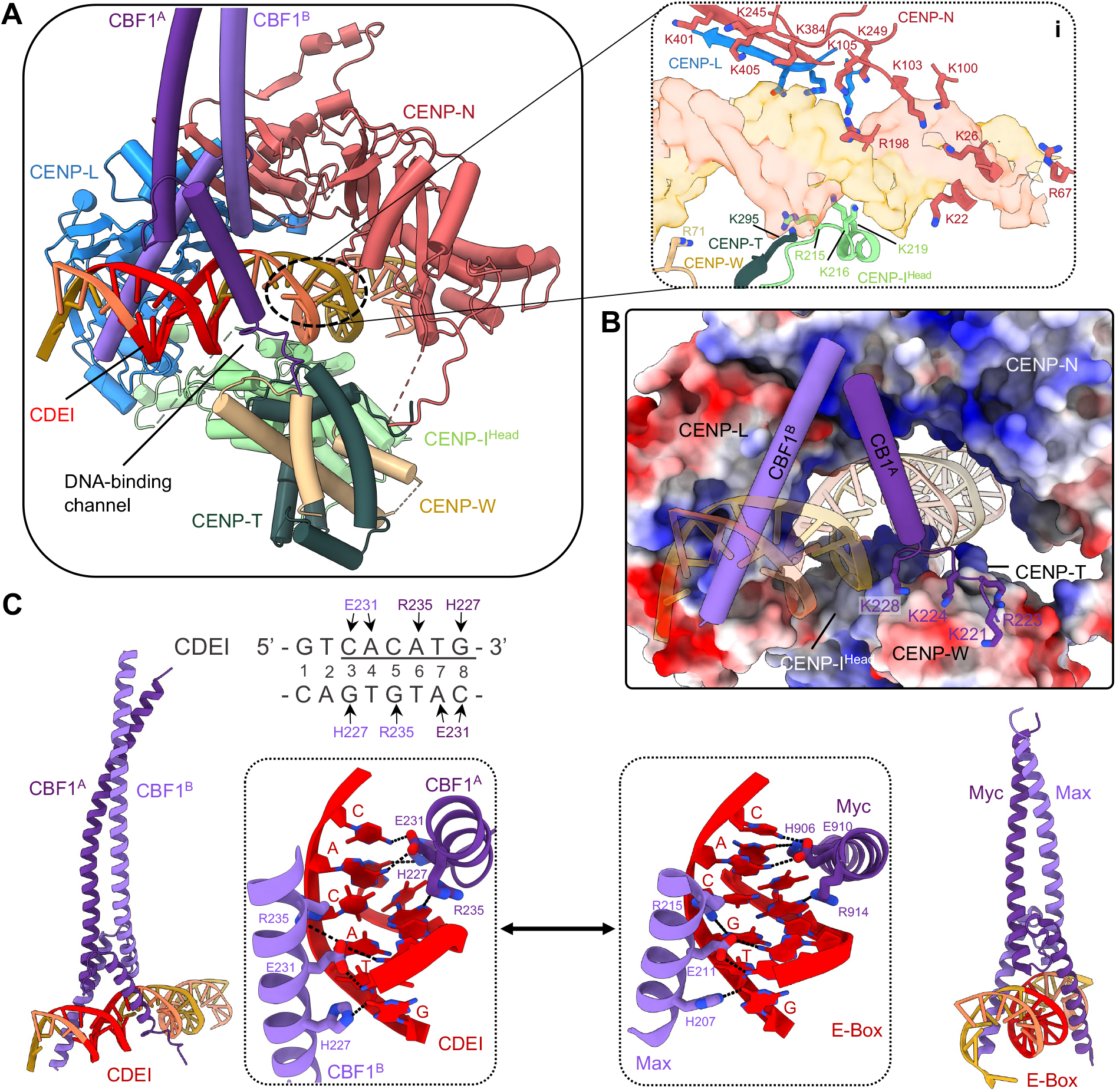
CBF1 extends the basic DNA-binding CENP-LN channel through interactions with CDEI. (**A**) Two subunits of CBF1 interact through their basic α-helices with the major groove of DNA. Inset i: CENP-LN, CENP-I^Head^ and CENP-TW form a basic, closed chamber that topologically entraps DNA. (**B**) The DNA-binding tunnel of CCAN has a marked electropositive potential and is extended by the basic helices of CBF1. An acidic patch on CENP-TW binds basic residues of the CBF1^A^ helix, thereby unfolding the N-terminal half of the helix. (**C**) The sequence-specific contacts of CBF1 with CAC(A/G)TG of CDEI are nearly identical to how the Myc-Max transcription factor interacts with its cognate E-box CACGTC motif.

We sought to assess the veracity of our CBF1:CCAN:DNA structure by testing the effects of mutants that disrupt either inter-subunit or CCAN:CENP-A^Nuc^ interactions, on the efficiency of mini chromosome segregation and sensitivity to microtubule poisons *in vivo*. Mutation of CDEIII and deletion of either CENP-N (*chl4Δ*) or CENP-I (c*tf3Δ*) severely compromised chromosome segregation efficiency (**Figures 3A-3C**). Mutations of basic residues of CENP-N that line the DNA-binding channel (*chl4*^*MT1*^) also severely impacted chromosome segregation efficiency (**Figures 3B and S4B**), consistent with our previous results that these mutations are synthetic lethal with a *cse4-R37A* mutant (a mutation within CENP-A^END^) (Yan et al., 2019). Disruption of basic residues of CENP-I^Head^ (*ctf3*^*MT1*^) that participate in the topological DNA-binding chamber (**Figures 2A and 2B**), significantly reduced chromosome segregation efficiency (**Figures 3C and S4B**). These results support our model that CBF1:CCAN engages a DNA duplex through the CENP-LN channel, augmented by contacts to CENP-I^Head^. It should be noted that the same basic patch on CENP-I^Head^ is in close proximity to the DNA gyre of CENP-A^Nuc^ in our previous CCAN:CENP-A^Nuc^ structure (Yan et al., 2019) (**Figures 1C and 1D**).

**Figure 3.**
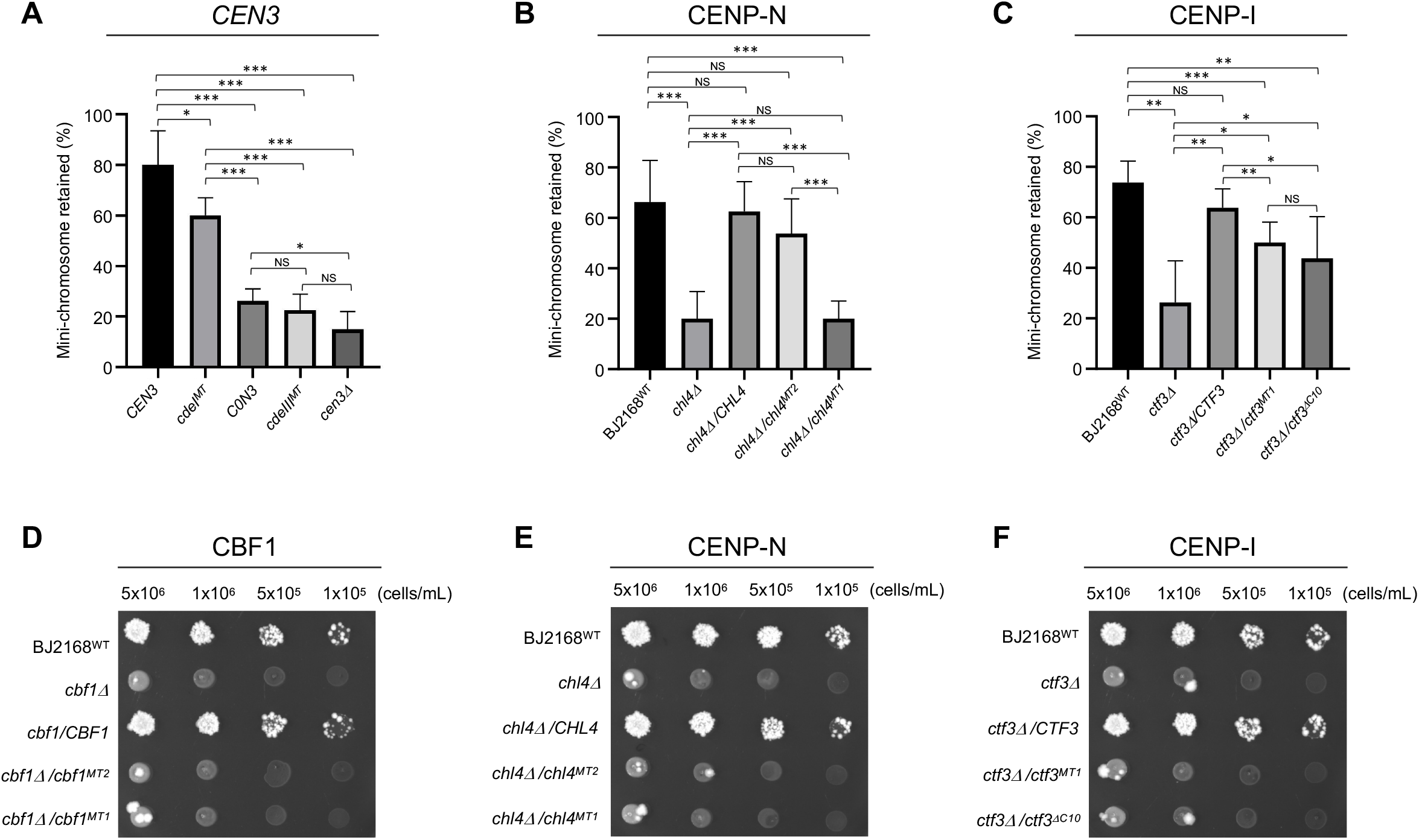
*In vivo* testing of the inner kinetochore structure. (**A-C**) Quantification of chromosome segregation loss for (**A**) *CEN3* mutants. *CEN3*: mini-chromosome with wild type *CEN3*; *cdeI*^*MT*^: CDEI mutant - GTCACATG to AATTGGCT; *C0N3*: mini-chromosome with *CEN3* replaced with *C0N3*; *cdeIII*^*MT*^: CDEIII mutant - CCG to AGC (Gal4-DNA-binding motif of CBF3 (Hegemann et al., 1988; Ng and Carbon, 1987; Yan et al., 2018)). *cen3Δ*: mini-chromosome with *CEN3* deleted. * p=0.05, ** p=0.01, *** p=0.001. (**B**) *CHL4* (CENP-N) mutants: *chl4*^*MT1*^ (DNA-binding groove: *chl4*^*K22S,K26S,R67S,K100S,K103S,K105S,R198S,K217S,K245S,K249S,K384S,K401S,K403S*^), *chl4*^*MT2*^ (histone H2A-H2B-binding: *chl4*^*D48R,D50R,E56R,E63R*^). (**C**) *CTF3* (CENP-I) mutants: *ctf3*^*MT1*^ (DNA binding: *ctf3*^*R215S,K216S,K219S,R222S,K225S*^), *ctf3*^*ΔC10*^ (CBF3 binding: *ctf3*^*F719S,Δ724-733*^). (**D-F**) Benomyl sensitivity spot assays for yeast strains harboring mutations of, (**D**) *CBF1*: *cbf1*^*MT1*^ (CENP-QU-binding mutant: *cbf1*^*L283E,L287W*^), *cbf1*^*MT2*^ (DNA-binding: *cbf1*^*K224S/K228S/R234S/R235S/K256S*^), (**E**) CHL4 (CENP-N): *chl4*^*MT1*^ and *chl4*^*MT2*^, and (**F**) CTF3 (CENP-I): *ctf3*^*MT1*^ and *ctf3*^*ΔC10*^, show sensitivity to benomyl.

The CBF1 homo-dimer interacts with the back-face of CCAN through its leucine zipper coiled-coil (residues 256-323) (**Figure S4A)**, forming a hydrophobic interface with CENP-Q, centered on CENP-Q^Ile292^ (**Figure S4A, inset ii**). This agrees with the observation that truncation and deletion of the leucine zipper disrupts both DNA binding by CBF1, and centromere function (Dowell et al., 1992). We assessed the consequence of disrupting this CBF1:CENP-QU interface (**Figure S4A, inset ii**). Replacing CBF1 residues Leu283 and Leu287, that form the CBF1:CENP-QU interface, with Glu and Trp, respectively (*cbf1*^*MT1*^), generated sensitivity to the microtubule-destabilizing drug benomyl, a phenotype identical to a CBF1 deletion (**Figures 3D and S4B**). CBF1 then contacts CDEI through basic residues of the bHLH, with His227, Glu231 and Arg235 of both CBF1 basic α-helices recognizing the bases of the near-palindromic CDEI motif: gtCAC[A/G]TG, in a sequence-specific manner (**Figure 2C**). The pattern of interactions between CBF1 and specific bases of CDEI correlates closely with both CDEI sequence conservation, and the effects of mutating individual positions on rates of chromosome loss (reviewed in (Gordon et al., 2011)). Indeed, deletion of the CDEI motif (*cdeI*^*MT*^) resulted in a significant mini-chromosome loss (**Figure 3A**). Strikingly, CBF1 binds to the CDEI motif in a manner that is identical to E-box (CACGTG) recognition by the heterodimeric Myc-Max transcription factor, as well as other dimeric Myc/Max/Mad bHLH factors, using the same conserved amino acid triplet (His, Glu, Arg) as CBF1 (**Figure 2C**) (Ferre-D’Amare et al., 1993; Nair and Burley, 2003). Mutation of either CBF1 residues mediating base-specific interactions, such as Glu231, or CBF1-binding nucleotides of CDEI, disrupted CDEI-CBF1 interactions (Masison et al., 1993; Mellor et al., 1990), and resulted in chromosome instability, and hypersensitivity to the mitotic drug thiabendazole (Foreman and Davis, 1993). To test the role of basic residues of the CBF1 bHLH motif, we replaced the wild type *CBF1* gene with a *cbf1* mutant in which DNA-binding residues were replaced with serines (*cbf1*^*MT2*^), and assessed the growth sensitivity of the mutant yeast strain to benomyl (Hyland et al., 1999). Whereas wild type *CBF1* rescued the benomyl sensitivity of a *cbf1Δ* strain, *cbf1*^*MT2*^ did not (**Figures 3D and S4B**). Lastly, the CBF1 basic helices adopt an asymmetric dimer conformation to accommodate the CCAN structure, most apparent for subunit CBF1^A^ where the N-terminal basic α-helix is unwound by nearly three turns to mediate contacts to CENP-TW (**Figure 2B**). Interestingly, this causes CBF1^A^ to form fewer DNA-backbone contacts to the right half-site of the CDEI palindrome (CAC[A/G]TG), perhaps explaining how mutations in this half-site affect centromere activity less than corresponding symmetrical changes in the left half-site (Niedenthal et al., 1991). In addition, the more complete α-helix of subunit CBF1^B^ interacts with CENP-L (**Figure S4A, inset iv**), an interface that might be disrupted when the left half-site of the palindrome is mutated, further rationalizing the asymmetric CDEI-disruption phenotype (Niedenthal et al., 1991).

### CBF1:CCAN topologically entraps DNA from the 5’ unwrapped end of *153C0N3*-CENP-A^Nuc^

During cryo-EM data processing we observed 2D-class averages, confirmed by a low-resolution 3D reconstruction, indicative of CCAN associated with *153C0N3*-CENP-A^Nuc^ (**Figure 4A**). We interpreted the small number of particles representing this reconstruction, together with its limited resolution, as evidence of sample denaturation during vitrification, despite an intact assembly in solution (**Figures S1B, S1C and S1D**). To stabilize *153C0N3*-CENP-A^Nuc^ for cryo-EM analysis we took advantage of the *CEN3*-CENP-A^Nuc^-stabilizing scFv fragment (Guan et al., 2021), and reduced sample complexity by omitting the CBF3^Core^ complex for the first iteration (**Figures S1E and S1F**). Because the binding site for scFv on H2A-H2B (Guan et al., 2021) overlaps with the CENP-C-binding site (Yan et al., 2019), we also omitted CENP-C from our reconstitution (CCAN^ΔC^). This approach enriched for stable CBF1:CCAN^ΔC^:*153C0N3*-CENP-A^Nuc^ on cryo-EM grids. A 3.4 Å consensus reconstruction was generated after refinement with an in-house program for refining flexible regions, specifically *153C0N3*-CENP-A^Nuc^ (**Figures S2B and S3B and Table S2**), and was identical to the low-resolution map reconstructed without scFv (**Figures 4A and 4B**). This structure (**Figure 4B**) showed that compared with the isolated CENP-A^Nuc^ reconstituted with either native *CEN3* (PDB 7K78, **Figure S5E**) (Guan et al., 2021) or Widom 601 DNA (PDB 70N1) (Migl et al., 2020), a further 10 bp of DNA are unwrapped from CENP-A^Nuc^ (a total of 33 bp at the 5’end). The unwrapped DNA is bound topologically in the CBF1:CCAN^ΔC^:CENP-A^Nuc^ complex, such that only ∼90 bp wrap the histone octamer. The dyad axis on *153C0N3*-CENP-A^Nuc^ aligns with the dyad axis on *CEN3*-CENP-A^Nuc^ (Guan et al., 2021), indicating that despite most of the *CEN3* CDEII being replaced by Widom 601 sequence in *153C0N3* (**Figures 1A and S1A**) the CENP-A octamer wraps the *153C0N3* and *CEN3* DNA sequences with identical positions. The CENP-A^Nuc^ is rotated relative to CCAN compared to our previous structure (Yan et al., 2019) (**Figures 1D and 4B**), reminiscent of human CCAN:CENP-A^Nuc^ (Yatskevich et al., 2022).

**Figure 4.**
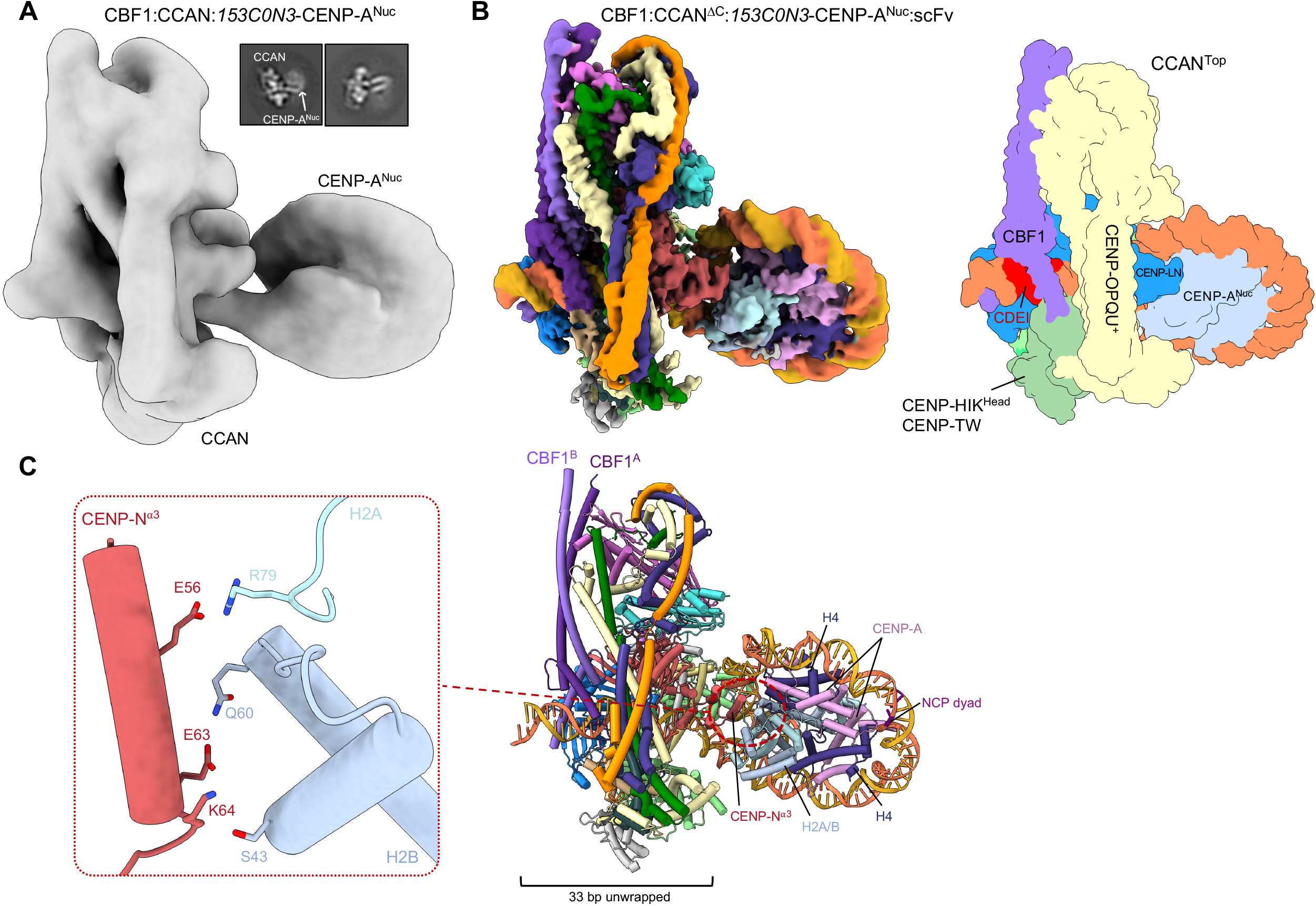
Structure of the CBF1:CCAN:CENP-A^Nuc^ complex. (**A**) Cryo-EM map of CBF1:CCAN:*153C0N3*-CENP-A^Nuc^ (no scFv) at 12 Å resolution. 11,435 particles were used for this reconstruction (0.75% of initial particles extracted). (**B**) Cryo-EM map of CBF1:*153C0N3*-CCAN^ΔC^:CENP-A^Nuc^:scFv, i.e., stabilized in the presence of scFv, at 3.4 Å resolution, showing how CBF1:CCAN^Top^ topologically entraps the unwrapped 5’-end of *153C0N3*-CENP-A^Nuc^. Schematic on the right. (**C**) Details of how CENP-N α3 helix interacts with basic residues of histones H2A-H2B that are exposed due to unwrapping of the CENP-A^Nuc^ DNA gyre. As such 33 bp of *153C0N3*-CENP-A^Nuc^ are unwrapped at its 5’-end.

### The inner kinetochore comprises two CCAN protomers bound to CENP-A^Nuc^ organized by the CBF1 and CBF3 complexes

Extending the scFv-stabilizing approach, we reconstituted the inner kinetochore complex by including CBF3^Core^ (**Figures S1G and S1I**). This CBF1:CCAN^ΔC^:CENP-A^Nuc^:CBF3^Core^:scFv complex will be referred to as the inner kinetochore (IK^*C0N3*^) from here onwards. Cryo-EM 2D-class averages revealed a much larger complex than in the absence of scFv, with CENP-A^Nuc^ clearly visible (compare **Figures S2C and 4A**). Consensus 3D reconstructions of the complex were refined to a limited 5.6 Å resolution due to conformational heterogeneity (**Figure S3C and Table S2**). Multibody refinement of rigid domains extended the resolution to 3.7-3.8 Å for all domains (**Figures 5A, 5B, S3C and S3D**). In this reconstruction we observed two CCAN protomers (as expected from SEC-MALS (**Figure S1D**)), *153C0N3*-CENP-A^Nuc^, a CBF1 homodimer, CBF3^Core^, and one scFv (**Figures 5 and S2D and Video S1**).

**Figure 5.**
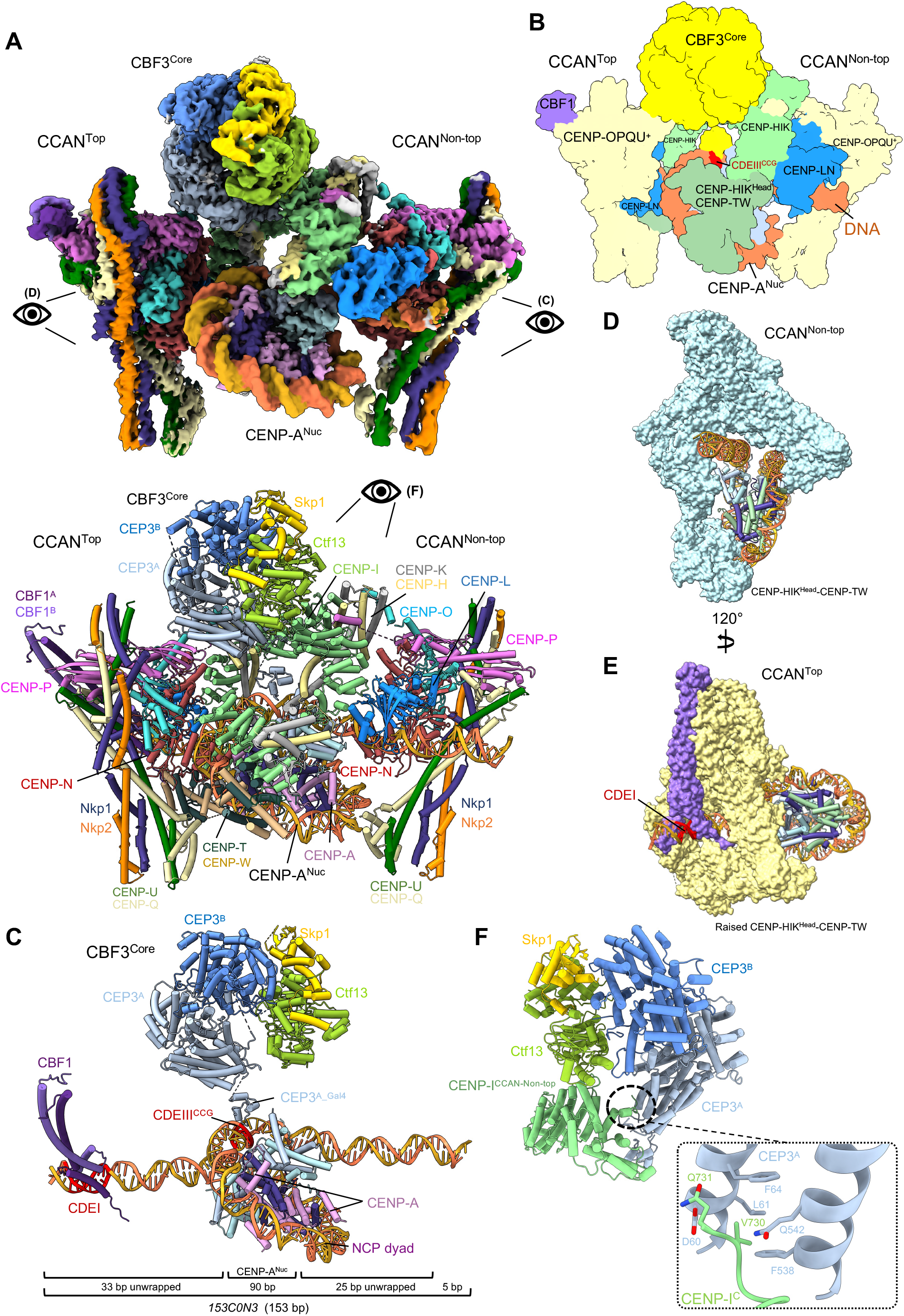
The inner kinetochore comprises two CCAN protomers bound to a central CENP-A^Nuc^ organized by the CBF1 and CBF3 complexes. (**A**) Cryo-EM map of the complex (top), and annotated ribbons representation (below). Two CCAN protomers flank the central CENP-A^Nuc^ in an asymmetric arrangement. CCAN^Non-top^ engages the 3’-end of the *153C0N3* DNA in an open configuration, identical to (Yan et al., 2019). CCAN^Top^ engages the 5’-end of the *153C0N3* and topologically entraps the DNA together with CBFI. CBF3^Core^ bridges the two CCAN modules. The single scFv bound to CENP-A^Nuc^ is not visible in this view (shown in Figure S2D). (**B**) Schematic of the complex. (**C**) CBF1 and CBF3^Core^ interact with CDEI and CDEIII, respectively. CDEI is located within the 5’ unwrapped DNA duplex of CENP-A^Nuc^, whereas CDEIII is located within the CENP-A^Nuc^ DNA gyre (SHL4). The body of CBF3^Core^ is distal to the face of the CENP-A nucleosome, a configuration that is different from (Guan et al., 2021), where the CBF3^Core^ sits proximal to the nucleosome (Figures S5A and S5B). The 153 bp *153C0N3* DNA is indicated: A total of 90 bp of DNA wraps the CENP-A^Nuc^ gyre, with 33 bp and 25 bp unwrapped at the 5’ and 3’ ends, respectively. (**D**) and (**E**) Views of CCAN^Non-top^ (D) and CBF1:CCAN^Top^ (E), in surface representation showing their different modes of binding CENP-A^Nuc^. (**F**) CEP3^A^ of CBF3 forms extensive contacts with the C-terminal region of CENP-I of CCAN^Non-top^. (**Video S1**).

In the inner kinetochore complex, CENP-A^Nuc^ is wrapped by only one turn of DNA (∼90 bp), in a left-handed configuration (**Figures 5A-C**). Both the 5’ and 3’ DNA ends of CENP-A^Nuc^ are therefore unwrapped, although to different extents: 33 and 25 bp at the 5’ and 3’ends, respectively (**Figures 5C, S5E and S5F**). The two unwrapped DNA duplexes create binding sites for two CCAN protomers (termed CCAN^Top^ and CCAN^Non-top^). These are arranged asymmetrically flanking the central CENP-A^Nuc^ (**Figures 5A and 5B**). CCAN^Top^ and CCAN^Non-top^ bind CENP-A^Nuc^ through the two different binding modes described above (i.e., both are observed in complexes with CENP-A^Nuc^ without scFv) (**Figures 5D and 5E**). CCAN^Top^, the protomer in complex with CBF1, is positioned at the 5’end of *153C0N3* with CBF1 binding the CDEI motif. CCAN^Top^ topologically entraps DNA through its CENP-LN channel engaging the unwrapped DNA duplex, and with the CENP-HIK^Head^-TW module positioned below, creates an enclosed DNA-binding chamber. This organization is identical to that observed in the CBF1:CCAN:DNA and CBF1:CCAN:*153C0N3*-CENP-A^Nuc^ complexes (**Figures 1B, 4A, 4B and 5E**). As was also seen for the CBF1:CCAN:*153C0N3*-CENP-A^Nuc^ complex, a 33 bp DNA duplex from the 5’end of *153C0N3* is unwrapped from CENP-A^Nuc^. Such a high degree of nucleosome unwrapping exposes basic residues of a H2A-H2B dimer responsible for binding DNA in a canonical H3 nucleosome (White et al., 2001). The removal of DNA-phosphate interactions from basic residues of H2A-H2B is partly compensated for by CCAN^Top^ through acidic residues on the α3 helix of CENP-N, a feature that is shared with the CBF1:*153C0N3*-CCAN:CENP-A^Nuc^ structure (**Figure 4C**). Mutating these residues of the CENP-N α3-helix (*chl4*^*MT2*^), did not impair chromosome segregation efficiency significantly (**Figures 3B and S4B**). However, yeast strains with this CENP-N mutant showed increased sensitivity to benomyl (**Figures 3E and S4B**), suggesting that loss of these CENP-N:H2A-H2B interactions causes a mildly deleterious effect on kinetochore stability.

The binding mode of CCAN^Non-top^, which has no associated CBF1, onto *153C0N3*-CENP-A^Nuc^, is the same as the CCAN assembled onto *W601*-CENP-A^Nuc^ (Yan et al., 2019) (**Figures 1F and 5D**). Similar to CCAN^Top^, CCAN^Non-top^ engages the unwrapped DNA duplex (in this case the 3’end) of *153C0N3*-CENP-A^Nuc^ (which lacks a CDEI motif) through its CENP-LN DNA-binding channel. However, *153C0N3*-CENP-A^Nuc^ is orientated differently in the CENP-LN channels of CCAN^Top^ and CCAN^Non-top^. For CCAN^Non-top^, CENP-A^Nuc^ engages the Y-shaped opening of the complex end on, such that the histone octamer lies below the CENP-LN channel (**Figure 5D**) (as for CCAN:*W601*-CENP-A^Nuc^ (**Figures 1D and 1F**) (Yan et al., 2019)). For CCAN^Non-top^, CENP-A^Nuc^ sterically obstructs the raised conformation of CENP-HIK^Head^-TW (**Figure 5D**). In contrast, for CCAN^Top^, CENP-A^Nuc^ is rotated by ∼150° about the unwrapped DNA duplex, bringing the histone octamer closer to CENP-LN. This orientation of CENP-A^Nuc^ requires further unwrapping of the DNA gyre to avoid CENP-A^Nuc^ clashing with CENP-LN, and it also allows space for the CENP-HIK^Head^-TW module to adopt a raised conformation below the CENP-LN channel, thereby generating the DNA chamber that topologically entraps the unwrapped DNA.

Together, the two CCAN protomers form an arch-like structure around CENP-A^Nuc^, bridged by CBF3^Core^. CBF3^Core^ interacts with CCAN^Non-top^ and CCAN^Top^ mainly through their CENP-I subunits (**Figures 5A and 5B**). Specifically, the C-terminus of CENP-I of CCAN^Non-top^ forms contacts with the CEP3^A^ subunit of CBF3^Core^, that are unique to CCAN^Non-top^ and not a feature of CCAN^Top^:CBF3 interactions (**Figure 5F**). Accordingly, deletion of the C-terminal 10 residues of CENP-I (*ctf3*^*ΔC10*^) caused severe chromosome segregation defects (**Figures 3C, 3F and S4B**). In the inner kinetochore complex, CBF3^Core^ adopts the same architecture as seen for free CBF3^Core^ (Leber et al., 2018; Lee et al., 2019; Yan et al., 2018; Zhang et al., 2018), except that the Gal4-DNA binding domain of the CEP3^A^ subunit shifts position to interact with the essential CCG motif of CDEIII **(Figure 5C**), similar to the complex of CBF3^Core^ with CENP-A^Nuc^ (Guan et al., 2021). The cryo-EM density for the CEP3^A^-Gal4 domain of CBF3^Core^ is diffuse, indicating that its interaction with the CDEIII motif as part of the inner kinetochore is weak (**Figure S5D**). A flexible linker connects the Gal4 domain to the globular domain of CEP3^A^ (**Figure 5C**). Relative to the CENP-A^Nuc^:CBF3^Core^ cryo-EM structure (Guan et al., 2021) (**Figure S5A**), in our inner kinetochore complex, the remainder of CBF3^Core^ is displaced from the face of CENP-A^Nuc^ (**Figure S5B**). This allows space for CENP-HIK-CENP-TW of CCAN^Non-top^, which in turn occupies the scFv-binding site on that face of CENP-A^Nuc^.

The bridging of the two CCAN protomers by CBF3^Core^ (**Figures 5A and 5B**) would suggest CBF3^Core^ contributes to both the stability and organization of the assembled inner kinetochore complex. Consistent with this is our observation that the cryo-EM 3D reconstruction of a complex comprising CBF1, CCAN^ΔC^, *153C0N3*-CENP-A^Nuc^ and scFv, described above (i.e., in the absence of CBF3^Core^), comprised predominantly a monomeric CCAN:CENP-A^Nuc^ assembly (**Figures 4B, S2B, S3B and S6A, column ii**). This cryo-EM dataset also produced 2D class averages with lower particle occupancy corresponding to a novel pseudo-symmetric di-CCAN:CENP-A^Nuc^ species, where the two CCAN protomers adopt the CCAN^Non-top^ configuration (**Figures S6A, column iii and S6D**). For the inner kinetochore sample in contrast, which includes CBF3^Core^, although we also observe this pseudo-symmetric di-CCAN species (with no bridging CBF3^Core^) (**Figure S6B, column iii**), the asymmetric di-CCAN of the inner kinetochore reconstruction (**Figure 5**) is substantially enriched, and is the predominant species (**Figure S6B, column iv**), consistent with the organizing role of CBF3^Core^.

### The inner kinetochore can be reconstituted with native *CEN3* DNA

We observed that the *153C0N3* DNA used in the *153C0N3*-CENP-A^Nuc^ reconstitution, in which the Widom 601 sequence partially substitutes for CDEII of *CEN3*, is positioned identically to the *CEN3* sequence of *CEN3*-CENP-A^Nuc^ (Guan et al., 2021) (**Figures S5E and S5F**). Specifically, the CDEI and CDEIII motifs in *153C0N3*-CENP-A^Nuc^ have the same position relative to the nucleosome dyad as their counterparts in *CEN3*-CENP-A^Nuc^. *153C0N3*-CENP-A^Nuc^ should therefore represent an effective and stable substitute of *CEN3*-CENP-A^Nuc^ for cryo-EM studies. Previous studies, however, had indicated that mutating CDEII results in mitotic delay and chromosome segregation defects (Baker and Rogers, 2005; Cumberledge and Carbon, 1987; Gaudet and Fitzgerald-Hayes, 1987, 1989; Murphy et al., 1991; Spencer and Hieter, 1992). We made similar observations in a chromosome segregation loss assay. A plasmid with a centromere based on *C0N3* is substantially more prone to mini-chromosome mis-segregation than a native *CEN3*-based plasmid, albeit better than an acentromeric plasmid *(cen3Δ*) (**Figure 3A**). This indicated that *C0N3* does not fully recapitulate *CEN3* function *in vivo*, possibly due to the lower nucleosome occupancy associated with W601 sequences *in vivo* (Perales et al., 2011). Therefore, we *in vitro* reconstituted the inner kinetochore complex using a fully native *153CEN3* DNA sequence (**Figures S1A and S1H**). As assessed by SEC, the *153CEN3* and *153C0N3* inner kinetochore complexes have similar compositions, eluting from the SEC column at identical volumes (**Figure S1I**). A cryo-EM data set indicated that the *CEN3* inner kinetochore is not stable on cryo-EM grids, despite scFv, most likely due to poor stability of *CEN3* nucleosomes, as also observed by others (Dechassa et al., 2011; Guan et al., 2021). Indeed, while we observed 2D classes with low particle occupancy for the pseudo-symmetric and asymmetric di-CCAN species (**Figure S6C, columns iii and iv**), the main species was a dimeric CCAN:DNA complex (**Figure S6A, column i**) (as described previously for *S. cerevisia*e apo-CCAN (Hinshaw and Harrison, 2019)), devoid of intact CENP-A^Nuc^.

### A CENP-A^Nuc^-CENP-QU pathway is independent of CCAN

Previous studies had identified an interaction between the essential N-terminal domain (END) of budding yeast CENP-A (residues 28-60: CENP-A^END^) (Chen et al., 2000; Keith et al., 1999) and two essential *S. cerevisiae* CCAN proteins, CENP-Q and CENP-U (Anedchenko et al., 2019; Fischbock-Halwachs et al., 2019). Reconstitution studies further showed that this CENP-QU pathway can transmit forces from the outer kinetochore to CENP-A^Nuc^ (Hamilton et al., 2020). Using ITC, we determined that a peptide modelled on CENP-A^END^ bound CENP-QU with a K_D_ ∼0.7 μM, consistent with an earlier study (Anedchenko et al., 2019) (**Figures S7A-S7E:** CENP-A^END-3^). Furthermore, residues 1-82 of CENP-A (CENP-A^N^) formed a stable complex with CENP-QU as assessed by SEC (**Figure S7F**). However, despite the stable CENP-QU:CENP-A^N^ association in solution, no CENP-A^N^ density was visible in the cryo-EM map of the CCAN:*W601*-CENP-A^Nuc^ complex (Yan et al., 2019) in the region of CENP-Q proposed to bind CENP-A^END^ (Anedchenko et al., 2019; Fischbock-Halwachs et al., 2019). To understand how CENP-A^N^ interacts with CENP-QU, we used AlphaFold2 (Jumper et al., 2021) to predict the structure of a CENP-A^N^:CENP-QU complex (**Figures 6A and S8A-S8C**). In the AlphaFold2 model, residues 21-80 of CENP-A form two α-helices arranged in a ‘V’ shape that embrace the four-helical bundle of CENP-QU^Foot^, and also contact the CENP-QU coiled-coil (**Figure 6A**). Consistent with prior CLMS data (Anedchenko et al., 2019; Fischbock-Halwachs et al., 2019), the majority of contacts involve CENP-Q. The longest of the two α-helices (residues 21 to 61) corresponds closely to the highly conserved CENP-A^END^ motif (**Figure 6A**) and its interaction at the composite CENP-QU interface is predicted with high confidence (**Figures S8A-S8C**). An extensive set of salt-bridge interactions define the CENP-A^N^:CENP-QU interface including CENP-A residues Arg37, Arg44, Arg46 and Lys49 that contact acidic residues on CENP-QU (**Figure 6A**). Mutation of these basic CENP-A residues was previously shown to cause chromosome segregation defects (Chen et al., 2000), whereas the interacting residues of CENP-QU were shown to bind CENP-A from mutagenesis and CLMS data (Anedchenko et al., 2019; Fischbock-Halwachs et al., 2019). In further support of our AlphaFold2 model, SEC analysis showed that substituting Lys residues for Glu191^CENP-U^, Glu194^CENP-U^ and Asp235^CENP-Q^, predicted to contact Arg37, Arg46 and Lys49 of CENP-A^N^ (CENP-QU^MT^) disrupted the CENP-A^N^:CENP-QU complex *in vitro* (**Figures 6A, S7F and S7G**).

**Figure 6.**
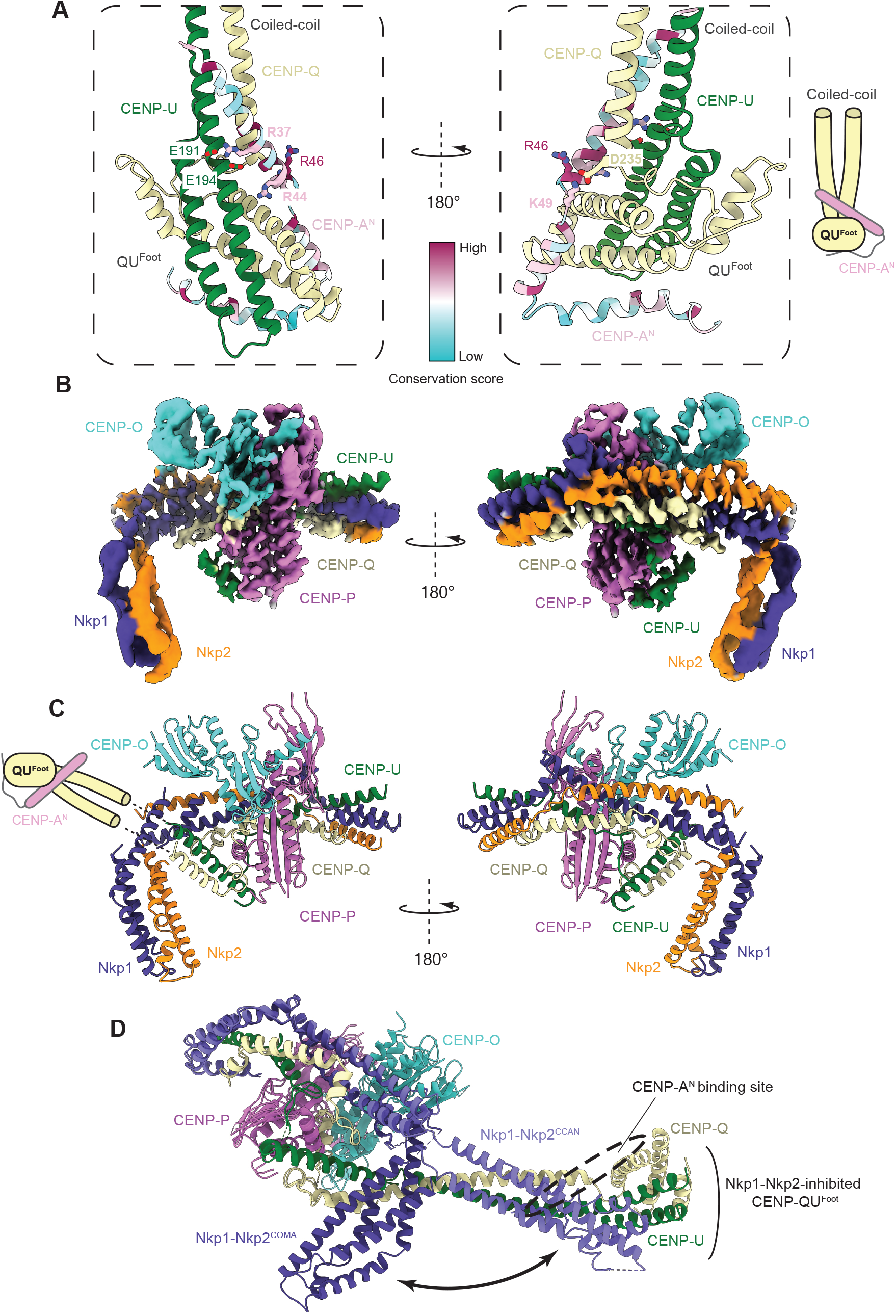
CENP-A^END^ interacts with CENP-QU auto-inhibited by CCAN. (**A**). Two views of an AlphaFold2 model predicting how CENP-A^N^ interacts with CENP-QU. The major site of interaction involves CENP-A^END^ with CENP-QU^Foot^ and CENP-QU coiled-coil. CENP-A^N^ is colored with a conservation score. (**B**) Cryo-EM map of the CENP-A^N^:CENP-OPQU+ complex. (**C**) Ribbons representation of the CENP-A^N^:CENP-OPQU+ complex. In this structure, CENP-QU^Foot^ and adjacent coiled-coil, including the CENP-A^END^-binding site of CENP-QU, are mobile and not visible in the cryo-EM map. Shown schematically. (**D**) Structure of CENP-OPQU+ in the context of CCAN superimposed onto the CENP-A^N^:CENP-OPQU+ complex. This shows the conformational change of the N-terminal region of Nkp1-Nkp2 exposing the CENP-A^END^-binding site on CENP-QU (mobile) in the free CENP-OPQU+ complex (**Video S2**). The *cse4-R37A* mutation causes a synthetic growth defect in cells lacking either CBF1 or CDE1 (Samel et al., 2012). Similar phenotypes are observed when the *cse4-R37A* mutant is combined with mutations in CCAN genes (Samel et al., 2012). Interestingly, a suppressor mutation of *cbf1Δ cse4-R37A* was identified in CENP-Q as a Cys substitution of Arg164 (*okp1-R164C*) (Anedchenko et al., 2019). In the CENP-QU:CENP-A^N^ model, Arg164^CENP-Q^ is at the CENP-QU:CENP-A^N^ interface, close to Arg46^CENP-A^ and Lys49^CENP-A^ (**A**). This close proximity of Arg164^CENP-Q^ to Arg46 ^CENP-A^ and Lys49^CENP-A^ might be expected to weaken CENP-QU:CENP-A^N^ interactions, and therefore, removal of a positive charge on CENP-Q in the *okp1-R164C* mutant would compensate for the loss of positive charge in CENP-A^N^ in the *cse4-R37A* mutant. Consistent with this prediction, co-immunoprecipitation studies showed that the *in vivo* association between Okp1 and Cse4-R73A is enhanced by the *okp1-R164C* mutant (Anedchenko et al., 2019). Furthermore, *in vitro*, a CENP-A^N^ peptide (residues 33-87) binds some 200-fold more tightly to CENP-Q^R164C^-CENP-U than to wildtype CENP-QU (Anedchenko et al., 2019).

Paradoxically, in the context of CCAN, the CENP-A^END^-binding site on CENP-QU is blocked by an N-terminal domain of the Nkp1-Nkp2 dimer, rendering CENP-QU incapable of binding CENP-A^END^ (**Figure 6D and Video S2**). Consistent with this, CENP-A^N^ did not bind either fully assembled CCAN or CENP-OPQU+ in the presence of CENP-LN (**Figures S9A and S9B**). CENP-LN likely promotes a conformational change in CENP-OPQU+, involving Nkp1-Nkp2, that blocks the CENP-A^END^-binding site on CENP-QU. To further assess this hypothesis, we determined a cryo-EM structure of the CENP-A^N^:CENP-OPQU+ complex to 3.4 Å resolution (**Figures 6B, 6C, S8D and S8E and Table S2 and Video S2**). Compared with its structure in the assembled CCAN, the free CENP-OPQU+ has a substantially altered conformation (**Figures 6B and 6C**). Specifically, the N-terminal domain of Nkp1-Nkp2 is bent back, contacting only the middle region of CENP-QU (**Figures 6B-6D**). The remodeled Nkp1-Nkp2 releases the coiled-coil domain of CENP-QU which is now able to bind CENP-A^END^. Although the CENP-A^END^-binding region of CENP-QU (CENP-QU^Foot^) is not visible in the cryo-EM map due to conformational heterogeneity, this structure, together with the Alphafold2 prediction, explains how CENP-A^END^ can bind CENP-QU in the presence of Nkp1-Nkp2, through release of CCAN-mediated auto-inhibition.

The CENP-A^END^-bound CENP-QU could represent a CCAN-independent axis of inner-outer kinetochore assembly. Consistent with this hypothesis, a CCAN:CENP-A^Nuc^ complex readily accommodated additional copies of CENP-QU, contingent on the N-terminal region of CENP-A (residues 1-129) (**Figures S9C and S9D**); and as described above, pre-assembled CCAN does not engage CENP-A^N^ (**Figure S9A**). Collectively, our results suggest that budding yeast CENP-A^Nuc^ recruits two fully assembled CCAN protomers that associate with CBF1 and CBF3, and can also bind two additional copies of CENP-QU (or the full CENP-OPQU+ complex). Our data are consistent with observations that CENP-QU is present at supernumerary amounts at kinetochores in cells, recruiting additional copies of the MIND and Ndc80 complexes (Dhatchinamoorthy et al., 2017). Lastly, *Cse4R37A* is a temperature-sensitive mutant (Samel et al., 2012) that, according to our structural model (**Figure 6A**), would specifically weaken the CENP-A^END^:CENP-QU interaction. *Cse4R37A* is synthetically lethal when combined with either deletion or mutation of other non-essential CCAN genes, such as CENP-N (Samel et al., 2012), that are crucial for CCAN assembly (Yan et al., 2019).

This further suggests that the CENP-A^N^:CENP-QU module and CCAN represent two distinct pathways to the outer kinetochore.

## Discussion

We present a structural model for the *S. cerevisiae* inner kinetochore reconstituted on an octameric CENP-A^Nuc^ which wraps one turn of left-handed helical DNA (**Video S1**). Substantial experimental studies previously demonstrated that native *S. cerevisiae* kinetochores consist of a single CENP-A^Nuc^ core (Furuyama and Biggins, 2007; Lang et al., 2018), comprising two CENP-A histones, (Aravamudhan et al., 2013; Camahort et al., 2009; Chen et al., 2000; Meluh et al., 1998; Wisniewski et al., 2014), that is perfectly positioned on centromeric sequences (Cole et al., 2011; Krassovsky et al., 2012). In our structure, the CENP-A^Nuc^ core is flanked on both sides by a pair of CCAN protomers that engage the unwrapped ends of the nucleosome asymmetrically. Two sequence-specific centromere-binding complexes, CBF1 and CBF3, function to organize the two CCAN protomers on CENP-A^Nuc^. The Ndc10 component of CBF3 only weakly associates with CBF3^Core^ (Guan et al., 2021), and its inclusion in our inner kinetochore reconstitution resulted in heterogeneous complexes. However, docking Ndc10 onto the inner kinetochore, guided by the CBF3^Holo^ structure (Yan et al., 2018), indicated a position that generates interfacial contacts with both CENP-I and CENP-L of CCAN^Non-top^ (**Figure S5C**).

The topological, CCAN^Top^-mediated entrapment of DNA by an enclosed chamber formed from CENP-LN:CENP-HIK^Head^-CENP-TW:CBF1 observed in our inner kinetochore complex reveals a mechanism of kinetochore attachment to centromeric chromatin that is evolutionarily conserved with how the human CCAN complex interacts with regional centromeres (Yatskevich et al., 2022). Differences, however, are that for budding yeast CCAN, the topologically entrapped DNA duplex is the unwrapped DNA terminus of CENP-A^Nuc^, whereas human CCAN interacts with the linker DNA of the 171 bp α-satellite repeats found at human centromeres. In yeast, the CDEI-targeting CBF1 component extends the basic DNA-binding channel and reinforces its topological closure through interactions with CENP-TW, in addition to its role in positioning CCAN^Top^ at the centromere. As in humans, yeast CENP-T and CENP-W have histone fold domains and, in vertebrate species, can tetramerize with CENP-SX to form nucleosome-like particles (Nishino et al., 2012; Schleiffer et al., 2012). In our reconstitutions and in previous work of others, CENP-SX did not assemble onto *S. cerevisiae* CCAN (Lang et al., 2018). Furthermore, the histone-like CENP-TW wrapping of centromere DNA at the human kinetochore (Yatskevich et al., 2022) is not conserved in budding yeast. Instead, CENP-TW of CCAN^Top^ forms few contacts with a weakly bent *CEN* DNA duplex. Our structure thus demonstrates a role for CENP-T and CENP-W together with CENP-I^Head^ and CBF1 in topologically entrapping centromeric DNA to bear spindle forces during mitosis.

Cryo-EM reconstructions of both *S. cerevisiae* and human CCAN revealed that their underlying architectures are highly conserved, including the central DNA-binding CENP-LN channel (Pesenti et al., 2022; Tian et al., 2022; Yatskevich et al., 2022). However, this CENP-LN channel is notably wider in *S. cerevisiae* CCAN compared with its counterpart in human CCAN. As noted by (Pesenti et al., 2022), based on AlphaFold2 predictions of CENP-LN from a variety of species, a wider channel appears conserved in yeast utilizing point centromeres, whereas the narrow CENP-LN channel is associated with organisms that evolved regional centromeres. Our structure of the *S. cerevisiae* inner kinetochore complex shows that only the wide CENP-LN channel is compatible with CCAN^Non-top^ engaging CENP-A^Nuc^ in an end-on, non-topological configuration. Other differences between the architectures of point and regional kinetochores include the stoichiometry of two CCAN protomers to CENP-A^Nuc^ in budding yeast, in contrast to a single human CCAN protomer associated with either one or two α-satellite repeat-CENP-A nucleosomes (Walstein et al., 2021; Yatskevich et al., 2022). Additionally, although the topological entrapment of DNA is conserved between human CCAN and the CBF1:CCAN^Top^ protomer of the *S. cerevisiae* inner kinetochore, the relative juxtapositions of CCAN and CENP-A^Nuc^ are not conserved.

Both CCAN protomers of the inner kinetochore assembly engage the unwrapped DNA ends of CENP-A^Nuc^. While the CCAN^Top^ protomer is positioned at CDEI, CCAN^Non-top^ extends 25 bp 3’ of CDEIII, so that the total length of DNA embedded in the inner kinetochore complex is ∼150 bp (**Figure 1A**). This matches almost exactly with the size and position of centromeric DNA protected from DNase1 and micrococcal nuclease digestion of native budding yeast chromosomes, including protection of the region extending ∼30 bp 3’ of CDEIII (Bloom and Carbon, 1982; Cole et al., 2011; Funk et al., 1989; Mellor et al., 1990; Wilmen and Hegemann, 1996). Thus, our structure is in good agreement with prior biochemical characterization of native *S. cerevisiae* kinetochore-centromere complexes, and is further supported by the functional roles of specific residues tested in the *in vivo* assays reported here.

Fourteen of the 16 budding yeast centromeres are between 117 and 119 bp in length, with *CEN4* being substantially shorter (111 bp), and *CEN12* the longest (120 bp) (Cole et al., 2011). Centromere length differences result from sequence variations in CDEII. As CDEII interacts with the dyad axis of the nucleosome, length variations in CDEII therefore predict that CDEI and CDEIII do not maintain the same position relative to the dyad axis in all 16 centromeres. Variations in CDEIII position relative to the dyad axis would require shifts in the Gal4-DNA binding domain of CBF3. We envision this is readily accommodated due to the flexible linker connecting the Gal4 domain to the main CEP3^A^ domain, and that the DNA gyre of CENP-A^Nuc^ is unobstructed by CCAN protomers in the immediate vicinity of CDEIII^CCG^ (**Figures 5A and 5C**). Variations in CDEI position relative to the CENP-A^Nuc^ dyad axis would involve a modest rotation of CENP-A^Nuc^ relative to CBF1:CCAN^Top^, possibly accommodated by the flexible interfaces connecting CCAN, CENP-A^Nuc^ and CBF3 components, as observed in our inner kinetochore cryo-EM reconstruction (**Figures S3C and S3D**).

By providing insight on the mechanism of CENP-A^END^ interactions with CENP-QU (**Figure 6**), we rationalize structural and genetic-dependency data by characterizing two independent CENP-A^Nuc^ pathways to the outer kinetochore. These involve (1) a direct link from CENP-A^END^ through CENP-U to MIND, independent of CCAN and (2) a CCAN:CENP-A^Nuc^ pathway linking CENP-U to MIND, and CENP-T to Ndc80 (**Figure 7**). CENP-A^END^ and CENP-N, as part of CCAN, function in separate pathways (1 and 2), potentially explaining how combined disruption of CENP-A^END^ and CENP-N causes synergistic effects (Samel et al., 2012). To which degree these pathways overlap physically at the centromere or whether they occur sequentially and transiently throughout mitosis remains unclear. It is possible, for example, that the CENP-QU pathway is also a centromere recruitment pathway for CCAN, and that CENP-A^END^ is displaced from CENP-QU^Foot^ by Nkp1-Nkp2 upon recruitment of CENP-LN and the other CCAN modules.

**Figure 7.**
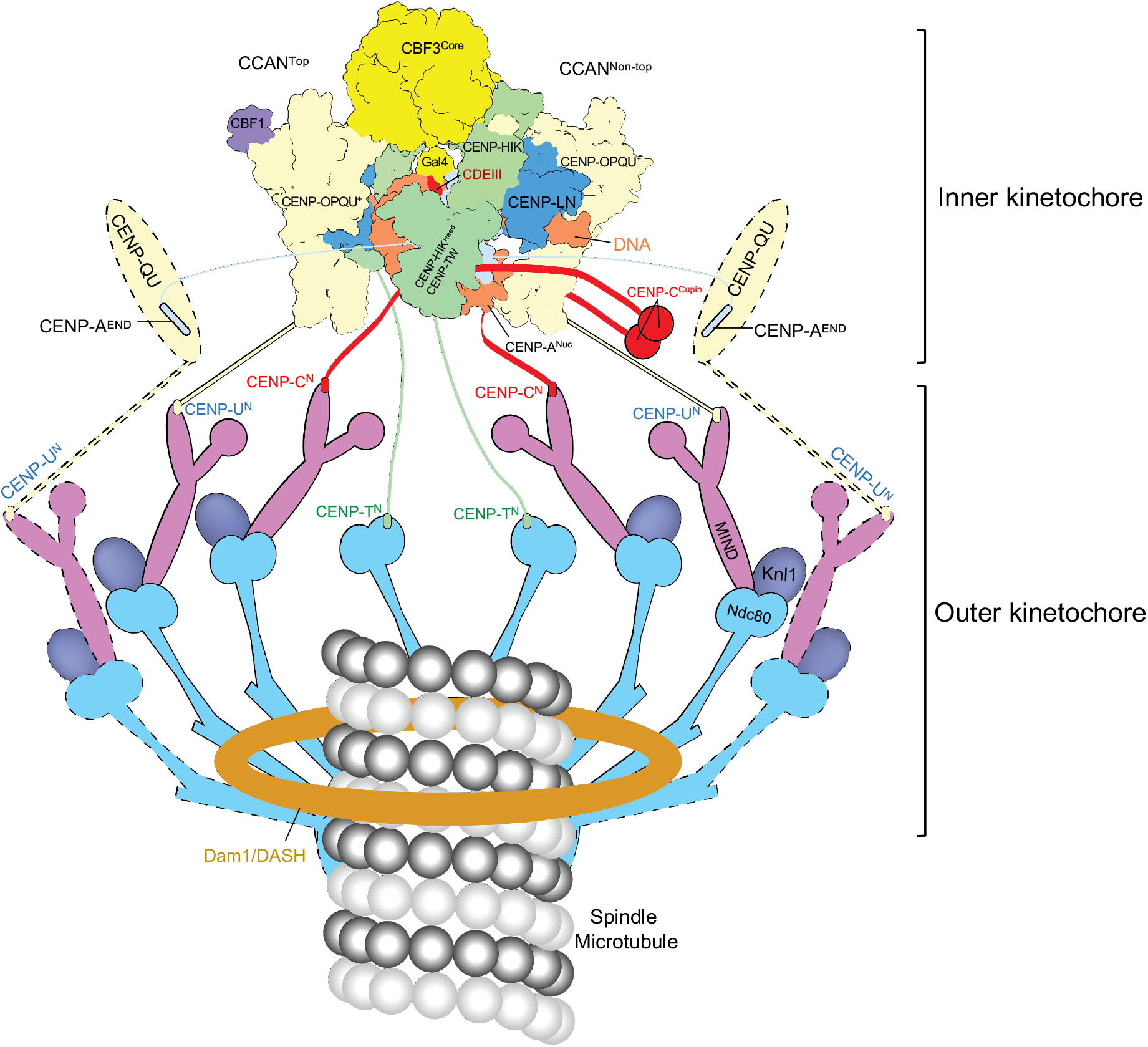
Schematic of the point centromere-kinetochore of *S. cerevisiae*. In this model the holo-inner kinetochore complex (two CCAN protomers) together with two CENP-A^END^-binding CENP-QU modules form a total of eight Ndc80 connections to the spindle microtubule. CENP-QU:MIND:Ndc80 modules with dashed outlines represent components of the CENP-QU pathway described in Figure 6. To which degree the CCAN and CENP-QU pathways overlap physically and temporally at the centromere remains to be understood. C^N^, U^N^, and T^N^ refer to the N-terminal motifs of CENP-C, CENP-U and CENP-T, respectively that bind MIND (C^N^, U^N^) and Ndc80 (T^N^).

The structural organization of inner kinetochore proteins observed here, that is the topological entrapment of DNA and CBF3-mediated bridging of the CCAN modules, underlies its function in force transduction from inner kinetochore to chromosome. This study demonstrates that topological entrapment of DNA is a kinetochore function conserved from yeast to humans (Yatskevich et al., 2022). What is lacking to construct a complete mechanistic model of kinetochore function is a molecular understanding of inner-outer kinetochore linkages at the centromere and how the outer kinetochore in turn tracks depolymerizing microtubule plus-ends. However, our structure now provides a compelling estimate for the probable stoichiometry of the inner and outer kinetochore, by accounting for all known attachment points. In our inner kinetochore model, up to six attachment points for the MIND and Ndc80 outer kinetochore modules are presented: two CENP-C molecules (binding a MIND module each), two CENP-T molecules (binding a Ndc80 module each) and two CENP-QU (binding a MIND module each, (Hornung et al., 2014)) (**Figure 7**). Taking into account the CENP-A^Nuc^-CENP-QU pathway described above, and assuming additional QU-modules assemble onto the inner kinetochore to interact with CENP-A^END^, an additional two MIND connection points are generated for a total of six MIND molecules (**Figure 7**). Interestingly, fluorescence microscopy studies estimate the number of MIND and Ndc80 components per kinetochore at ∼6-7 and ∼8-10, respectively (Cieslinski et al., 2021; Joglekar et al., 2006). It should be noted, however, that the same studies report other CCAN components at either ∼one copy per kinetochore (Joglekar et al., 2006) or ∼2-4 copies per kinetochore (Cieslinski et al., 2021) depending on the fluorescence reference standard. FRAP and photoconversion experiments have shown that additional copies of Ndc80, MIND and CENP-PQU modules are incorporated at the centromere during anaphase, to form an ‘anaphase configuration’ kinetochore, which is postulated to reinforce the load-bearing attachment (Dhatchinamoorthy et al., 2017). Although speculative, it is possible that temporal dynamics of kinetochore composition and layout during the cell cycle, such as those described above, can account for yet unresolved observations such as the CENP-QU pathway.

In conclusion, our structure of the budding yeast inner kinetochore reconstituted onto a CENP-A nucleosome with near-native centromeric DNA provides a foundation for understanding the higher-order centromere-kinetochore assembly, with implications for the architecture of regional centromeres and with insights into KMN stoichiometry. This study answers long-standing questions of how the defined sequence elements of point centromeres recruit sequence-specific DNA-binding complexes to organize the load-bearing attachments of the inner kinetochore. A striking mode of DNA engagement by CCAN^Top^ involves topological entrapment within a DNA-binding chamber. Conservation of this architectural feature, from the point centromeres of budding yeast to the much larger regional centromeres of humans, suggests a highly robust and effective mechanism has evolved to generate the stable attachment of chromosomes to the mitotic spindle for the maintenance of chromosome stability

## Acknowledgements

We acknowledge Diamond Light Source for access and support of the cryo-EM facilities at the UK’s national Electron Bio-imaging Centre (eBIC) [under proposal EM BI31336], funded by the Wellcome Trust, MRC and BBRSC. We are grateful to the LMB EM facility for help with the EM data collection, to J. Grimmett, T. Darling and I. Clayson for high-performance computing, and to J. Shi for help with insect cell expression. We thank K. Muir and N. Turner for discussions and members of the DB group for comments on the manuscript. This study was supported by UKRI/Medical Research Council MC_UP_1201/6 (D.B.) and MC_UP_ A025_1013 (S.W.H.S.), Cancer Research UK C576/A14109 (D.B.), and a Boehringer Ingleheim Fonds Fellowship (S.Y.), J.S. is supported by the Alan Turing Institute (EPSRC Grant EP/W006022/1, in particular the “AI for Science” theme within that grant). For the purpose of open access, the author has applied a CC BY public copyright licence to any Author Accepted Manuscript version arising.

## Author contributions

T.D., Z.Z., J.Y., S.Y. and D.B. designed the study and experiments. Z.Z. cloned constructs and prepared nucleosomal DNA. J.Y. and Z.Z. purified proteins and nucleosomes. J.Y. assembled complexes for structure determination. T.D. and S.Y. prepared and optimized cryo-EM grids, collected and processed data and built structures. AlphaFold2 predictions were performed by S.Y. and D.B. Z.Z. devised and performed the genetic stability assays. S.H.M. performed SEC-MALS. D.B. performed the ITC experiments. J.Y. and Z.Z. performed biochemical assays. J.S. and S.H.W.S. devised and implemented the deformation refinement method. T.D., S.Y. and D.B. wrote the manuscript with input from others.

## STAR Methods

### Resource Availability

#### Lead contact

Further information and requests for resources and reagents should be directed to and will be fulfilled by the Lead Contact, David Barford.

#### Materials Availability

All unique/stable reagents generated in this study are available from the Lead Contact with a completed Materials Transfer Agreement.

#### Data and code availability

The cryoEM maps have been deposited to the Electron Microscopy Data Bank (EMDB) with accession numbers EMD-xxxx and EMD-xxxx. The protein model has been deposited to the Protein Data Bank (PDB) with accession number xxxx.

### Method Details

#### Peptide Synthesis

CENP-A^END^ peptides were synthesized by Cambridge Research Biochemicals at 95% purity. All three peptides, contained N-terminal acetylation, C-terminal amidation and were at 95% purity (W) = peptide with an additional N-terminal tryptophan was used to confirm peptide concentration in stoichiometry studies. CENP-A^END-1^: DA<u>SINDRALSLLQRTRAT</u>DAW residues 32-48; CENP-A^END-2^: <u>AGDQQSINDRALSLLQRTRATKN</u>W residues 28-50; CENP-A^END-3^: <u>AGDQQSINDRALSLLQRTRATKNLFPRREERRR</u>W residues 28-60.

#### ITC

Isothermal titration calorimetry (ITC) was performed using an Auto-iTC200 instrument (Malvern Instruments, Malvern, UK) at 20°C. CENP-A^END-1^, CENP-A^END-2^, CENP-A^END-3^: (Kd of 11.5, 1.0, 0.72 μM). Peptide concentrations (1): 2.18 mM, (2): 1.82 mM, (3): 1.18 and 1.24 mM. Protein concentration in well: 0.142 mM and 0.18 mM. Buffer: 20 mM Hepes (pH 7.5), 100 mM NaCl, 1 mM TCEP. For each titration run, 370 μL of CENP-QU (between 142-180 μM) was used to load the calorimeter cell. The CENP-A^END^ peptides at 1.18-2.18 mM were titrated into the cell consisting of one 0.5 μl injection followed by 19 injections of 2 μl each. After discarding the initial injection, the changes in the heat released were integrated over the entire titration and fitted to a single-site binding model using the MicroCal PEAQ-ITC Analysis Software 1.0.0.1258 (Malvern Instruments). Titrations were performed in triplicate.

#### Cloning

All genes and proteins used in this study are of *S. cerevisiae* origin. Expression constructs and systems for assembly of the CENP-OPQU^+^, CENP-HIKTW and CENP-LN complexes, CENP-C and the CENP-A octamer were described in (Yan et al., 2019) (**Table S1**). For CENP-A^ΔN^ octamer preparation, the expression cassette of CENP-A^130-229^ was combined with *H2A, H2B* and *H4* expression cassettes in a single pET28 plasmid. The CBF3^Holo^ complex was prepared as described in (Yan et al., 2018). The CBF3^Core^ complex (*Cep3, Ctf13* and *Skp1*) was cloned into pU2, and Cbf1 was cloned into pU1 (Zhang et al., 2016). Cep3 and Cbf1 were cloned with C-terminal TEV-cleavable double StrepII tags as described in (Zhang et al., 2016). For the CENP-ΔQU complex, the coding sequences of CENP-Q^1-294^ (CENP-ΔQ) and CENP-U^30-266^ (CENP-ΔU) were cloned into pET28 plasmids, with a TEV cleavable double StrepII tag on the CENP-U C-terminus. The two cassettes were further combined into a single pET28 plasmid. For CENP-A^N^ protein, the coding region of CENP-A^1-82^ was cloned into pAcycDuet plasmid with a N-terminal 3C protease cleavable His_6_ tag.

For the single chain antibody, the scFv coding sequence (Guan et al., 2021) was synthesized (Thermo Fisher Scientific) and sub-cloned into pET28A.

#### Protein and complex preparation

CCAN subcomplexes (CENP-C, CENP-LN, CENP-OPQU+ and CENP-HIKTW) were expressed in the insect cell-baculovirus system, CENP-A and CENP-A^ΔN^ octamers in *E. coli*, and purified as described in (Yan et al., 2019). CBF1, CBF3^Core^, CBF3^Holo^ and Ndc10 were expressed individually in High-5 insect cells. Cells were harvested 48 h after infection. The cleared lysate was loaded onto an affinity column (either Strep-Tactin column (Qiagen) or HisTrap HP column (Qiagen)) for purification of expressed proteins and sub-complexes. The tags were cleaved by a 16 h incubation with TEV protease at 4°C. The protein complexes were then purified by Resource Q anion-exchange, and further purified by size-exclusion chromatography in a buffer of 20 mM Hepes (pH 8.0), 300 mM NaCl, 0.5 mM TCEP.

CENP-ΔQU complex expression was performed at 20°C for 16 h with 0.36 mM IPTG in *E. coli* strain B834 with codon plus Rare2. The complex was purified by a combination of Strep-Tactin (Qiagen) in a buffer of 50 mM Tris.HCl (pH 8.0), 250 mM NaCl, 1 mM EDTA and 1 mM DTT, followed by cation exchange chromatography Resource S (Cytiva) with buffer of 20 mM Tris.HCl (pH 7.5), 75 mM NaCl, 1 mM EDTA and 1 mM DTT (gradient elution with buffer containing 1 M NaCl), and Superdex 200 size-exclusion chromatography (Cytiva) with buffer of 10 mM Tris.HCl (pH 7.5) 150 mM NaCl, 1 mM DTT and 1 mM EDTA. The complex was concentrated to 8 mg/mL and stored at -80°C.

CENP-A^N^ was expressed as for CENP-QU. The protein was purified by Ni-NTA with buffer of 50 mM Tris.HCl (pH 8.0), 250 mM NaCl, eluted with the buffer containing 300 mM imidazole. The CENP-A^N^ was further separated by Superdex 75 size-exclusion chromatography (Cytiva) with buffer of 10 mM Tris.HCl (pH 7.5), 150 mM NaCl, 1 mM DTT and 1 mM EDTA. The protein was concentrated to 2 mg/mL and stored at -80°C The single chain antibody fragment (scFv) was prepared using a protocol adapted from (Guan et al., 2021). The inclusion bodies that contain scFv were prepared from overexpressing scFv from a pET28A plasmid in *E. coli* B834^rare2^ cells. The inclusion bodies were solubilized with a denaturation buffer of 100 mM Tris.HCl (pH 8.0), 6 M guanidine buffer, 2 mM EDTA, and the spun down. The supernatant was adjusted to a protein concentration to 10 mg/mL. 1,4-dithioerythritol (DTE) powder was added to a final concentration of 10 mg/mL and shaked at 20°C for 16 h. While stirring, 10 mL of the supernatant was quickly added to 1 liter of pre-chilled (10°C) refolding buffer and stirred for 3 mins. The refolding buffer contains 551 mg/L freshly add oxidized glutathione powder in 100 mM Tris.HCl (pH 9.5), 1 mM EDTA, 0.5 M arginine, pH 9.5. The refolding solution was incubated at 10°C for 48 h without stirring. 1 L of refolding solution was then dialyzed against 5 L of pre-chilled (4°C) dialysis buffer of 20 mM Tris.HCl (pH 7.4) with 34 g of urea (added before dialysis) with a 6-8 kDa cut-off dialysis tubing for 16 h at 4°C. This dialysis step was then repeated using fresh buffer.

The refolding solution was filtered through a 0.22 μM filter unit then mixed with 4 mL of pre-equilibrated SP Sepharose Fast Flow resin in SP-binding buffer of 20 mM Tris.HCl (pH 7.4) for 1 h at 4°C. The resin was collected with an Econo column, washed with SP-binding buffer and the scFv was eluted using 360 mM NaCl in the SP-binding buffer. The scFv was further purified on a Superdex S75 size-exclusion column, concentrated to 1 mg/mL and stored at -80°C in a buffer of 20 mM Tris.HCl (pH 7.4), 150 mM NaCl, 1 mM EDTA.

#### DNA generation

*153C0N3* DNA fragment was prepared by primer-extension method. Oligos of C0N3F ATAAGTCACA TGGTGCCGAG GCCGCTCAAT TGGTCGTAGA CAGCTCTAGC ACCGCTTAAA CGCACGTA CG CGCTGTCCCC CGCG TTTTAA and C0N3R TTCAATGAAA TATATATTTC TTACTATTTC TTTTTTAACT TTCGGAAATC AAATACACTA ATATTAAAAC GCGGGGGACA GCGCGTACGT were synthesized by Sigma-Aldrich. After mixing the oligos in 1x PCR reaction mixture, the fragment was produced with one step extension at 68°C for 1 min. The final product of 153 base pair *153C0N3* of ATAAGTCACA TGGTGCCGAG GCCGCTCAAT TGGTCGTAGA CAGCTCTAGC ACCGCTTAAA CGCACGTACG CGCTGTCCCC CGCGTTTTAA TATTAGTGTA TTTGATTTCC GAAAGTTAAA AAAGAAATAG TAAGAAATAT ATATTTCATT GAA fragment was purified using a 1 mL Resource Q anion exchange chromatography and stored in a buffer of 2 M NaCl, 10 mM Tris.HCl (pH 7.5), 1 mM EDTA, 2 mM DTT at -20°C.

For *153CEN3*, three copies of (ATAAGTCACA TGATGATATT TGATTTTATT ATATTTTTAA AAAAAGTAAA AAATAAAAAG TAGTTTATTT TTAAAAAATA AAATTTAAAA TATTAGTGTA TTTGATTTCC GAAAGTTAAA AAAGAAATAG TAAGAAATAT ATATTTCATT GAA) flanked by EcoRV site were cloned into pUC19. The plasmid was isolated by using the Plasmid Giga Kit (QIAGEN). The 153CEN3 fragment was purified with a 1 mL Resource Q anion exchange chromatography column (Cytiva) after digestion with EcoRV-HF (NEB) for 16 h. The purified DNA was precipitated, dissolved, buffer-exchanged and stored in a buffer of 2 M NaCl, 10 mM Tris.HCl (pH 7.5), 1 mM EDTA, 2 mM DTT at -20°C.

#### CENP-A nucleosome and derivatives preparation

CENP-A and CENP-A^ΔN^ nucleosomes were prepared by wrapping the prepared octamers with *153C0N3* DNA or *153CEN3* DNA by gradient dialysis. Either CENP-A or CENP-A^ΔN^ octamers were mixed with either *153C0N3* DNA or *153CEN3* DNA all at 7.8 μM. The mixture was dialyzed from 2 M NaCl to 100 mM NaCl in 10 mM Tris.HCl (pH 7.4), 1 mM EDTA, 2 mM DTT buffer for at least 16 h at 20°C. The mixture was further dialyzed in a buffer of 10 mM Tris.HCl (pH 7.4), 1 mM EDTA, 2 mM DTT for 4 h. For the *153CEN3*-CENP-A nucleosome, the final dialysis step was performed at 65°C for 4 h, then spun down for 1 min to remove aggregates at 4°C. The wrapped nucleosomes were assessed on native agarose gels and stored at 4°C.

#### Assembly of CBF1:CCAN^ΔC^:*153C0N3*-CENP-A^Nuc^:CBF3^Core^:scFv and CBF1:CCAN^ΔC^:*153CEN3*-CENP-A^Nuc^:CBF3^Core^:scFv complexes

CENP-A nucleosome was mixed with CCAN sub-complexes: CENP-LN, CENP-OPQU^+^, CENP-HIK-TW, CBF1 and CBF3^Core^ at 2 μM concentration. The mixture was dialyzed in a buffer of 20 mM Hepes (pH 8.0), 80 mM NaCl for at least 5 h to remove DTT or TCEP. scFv (4 μM) was then added, and the sample dialyzed against a buffer of 20 mM Hepes (pH 8.0), 50 mM NaCl for 14 h at 4°C. The complex was then concentrated to 3 mg/mL. To stabilize the complexes, 3 mM BS3 was used to cross-link the complex for 30 mins on ice. The reaction was quenched by 50 mM Tris.HCl (pH 8.0) and incubated on ice for 20 min. The mixture was applied to an Agilent 1000Å column to remove excess CCAN sub-complexes before preparing cryo-EM grids. Uncross-linked complex was also loaded on to an Agilent 1000Å column to access the quality of the assembled complex. The same procedure was applied for CBF1:CCAN:*153C0N3*-CENP-A^Nuc^, but without CBF3^Core^.

#### Assembly of CBF1:CCAN:*153C0N3*-CENP-A^Nuc^:CBF3^Core^

As for CBF1:CCAN^ΔC^:*153C0N3*-CENP-A^Nuc^:CBF3^Core^:scFv except that CENP-C was included with CCAN sub-complexes when mixed with CENP-A^Nuc^, and scFv was omitted.

#### Testing supernumerary CENP-QU binding to CCAN:CENP-A nucleosome complexes mediated through CENP-A^N^

CENP-A or CENP-A^ΔN^ nucleosomes were wrapped with *153C0N3* DNA. The nucleosomes were then mixed with CCAN components (CENP-C, CENP-LN, CENP-OPQU+ and CENP-HIK-TW) to form CCAN:*153C0N3*-CENP-A or *153C0N3* CENP-A^ΔN^ nucleosome complexes. CENP-ΔQU was mixed with either of the two complexes at 2 mM in a buffer of 20 mM Hepes (pH 8.0), 80 mM NaCl and 0.5 mM TCEP for 2 h. The mixtures were then loaded onto an Agilent 1000Å column. The peak fractions were visualized by 4-12% on an SDS-PAGE gel stained with Instant Blue Coomassie.

#### CENP-OPQU+:CENP-A^N^ sample preparation for cryo-EM

To generate CENP-OPQU+:CENP-A^N^ complexes, 10 μM of CENP-OPQU+ was incubated with 10

μM CENP-A^N^ in buffer of 20 mM Hepes (pH 8.0), 80 mM NaCl, 0.5 mM TCEP on ice for 1 h, and then loaded onto an Agilent 1000Å column. The eluted samples were visualized by SDS-PAGE stained with Instant Blue Coomassie. To prepare cryo-EM grids, the CENP-OPQU+:CENP-A^N^ complex was cross-linked by incubation in 3 mM BS3 on ice for 30 min, followed by quenching with 50 mM Tris.HCl (pH 8.0) on ice for 20 mins.

#### Assessment of CENP-A^N^ binding to CENP-OPQU+ in the presence of CENP-LN

To test the effect of CENP-LN on the CENP-OPQU+:CENP-A^N^, complex, CENP-A^N^, CENP-OPQU+ and CENP-LN were mixed at 4 μM each in a buffer of 20 mM Hepes (pH 8.0), 80 mM NaCl, 0.5 mM TCEP and loaded onto a Superose 6 size-exclusion column.

#### Test binding of CENP-A^N^ to CCAN

To test binding of CENP-A^N^ to CCAN, CENP-A^N^ (2.5 μM) and CCAN components (CENP-C, CENP-LN, CENP-OPQU+ and CENP-HIK-TW) (2.0 μM) were mixed in a buffer of 20 mM Hepes (pH 8.0), 80 mM NaCl, 0.5 mM TCEP and loaded onto an Agilent 1000 size-exclusion column.

#### SEC-multi-angle light scattering (SEC-MALS)

Size-exclusion chromatography coupled with multi-angle static light scattering (SEC-MALS), was performed using an Agilent 1200 series LC system with an online Dawn Helios ii system (Wyatt) equipped with a QELS+ module (Wyatt) and an Optilab rEX differential refractive index detector (Wyatt). CENP-A nucleosome and all the CCAN sub-complexes: CENP-C, CENP-LN, CENP-OPQU^+^, HIK-TW, together with CBF1 and CBF3^Core^ complexes were mixed at 2 μM concentration to generate the complete inner kinetochore assembly. The mixture was dialyzed in a buffer of 20 mM Hepes (pH 8.0), 80 mM NaCl, 0.5 mM TCEP for at least 5 h. The inner kinetochore sample was then cross-linked with 3 mM BS3 for 30 min. The cross-linked sample was purified on an Aligent 1000Å column. The peak fractions were concentrated and 100 μl was injected onto an Agilent Bio SEC-5 column gel filtration column pre-equilibrated in 10 mM Hepes (pH 8.0), 80 mM NaCl, 1 mM EDTA and 0.5 mM TCEP. The light scattering and protein concentration at each point across the peaks in the chromatograph were used to determine the absolute molecular mass from the intercept of the Debye plot using Zimm’s model as implemented in the ASTRA v7.3.0.11 software (Wyatt Technologies).

To determine inter-detector delay volumes, band-broadening constants and detector intensity normalization constants for the instrument, thyroglobulin was used as a standard prior-to sample measurement. Data were plotted with the program PRISM v8.2.0 (Graph Pad Software Inc.).

#### Minichromosomal stability assay

A fragment of *ARS1-TRP1-153CEN3* was cloned into the *pUC18* plasmid to generate a *153CEN3* mini-chromosome (wildtype: *CEN3*). Based on *CEN3, cdeIII*^MT^ was generated by exchanging CCG to AGC. *cdeI*^MT^ was created by exchanging its GTCACATG to AATTGGCT. The C0N3 mini-chromosome was generated by exchanging its *153CEN3* with *153C0N3*. The sequence of *153CEN3* was removed from *CEN3* for the *cen3Δ* mini-chromosome control. This set of mini-chromosomes was transformed into BJ2168, and selected with Sc-TRP (yeast synthetic medium drop out tryptophan) plates. A single colony from each was cultured in non-selective YPD medium for 12 h. The cultures were diluted and spread onto YPD plates and grown for 3 days to obtain single colonies. The colonies were then plated onto Sc-TRP plates and incubated for 3 days at 30 °C, and the selected colonies were counted to determine the percentage of mini-chromosome retained.

The BJ2168^*CEN3*^ strain was used for deletion of the *CBF1, CTF3* and *CHL4* genes by replacing their respective coding sequences with the KanMX6 gene to create the BJ2168^*153CEN3,cbf1Δ*^, BJ2168^*153CEN3,ctf3Δ*^ and BJ2168^*153CEN3,chl4Δ*^ strains by selection on G418 plates. The knockout strains were confirmed by sequencing.

*CBF1, CTF3* (CENP-I) and *CHL4* (CENP-N) genes were cloned into pYes2 plasmids along with their native promoters and the *URA3* selection marker. *cbf1*^*MT1*(L283E,L287W)^, *cbf1*^*MT2*(K224S,K228S,R234S,R235S,K256S^, *ctf3*^*MT1*(R215S,K216S,K219S,R222S,K225S)^, *ctf3*^ΔC10(F719S,Δ724-733)^, *chl4*^MT1(K22S,K26S,R67S,K100S,K103S,K105S,R198S,K217S,K245S,K249S,K384S,K401S,K403S^, *chl4*^MT2(D48R,D50R,E56R,E63R)^ mutants were created from their wildtype constructs. These plasmids were transformed into the appropriate BJ2168^*CEN3*^ knockout strain to create BJ2168^*CEN3,cbf1Δ,CBF1*^, BJ2168^*CEN3,cbf1Δ,cbf1-MT1*^, BJ2168^*CEN3,cbf1Δ,cbf1-MT2*^, BJ2168^*CEN3,ctf3Δ,CTF3*^, BJ2168^*CEN3,ctf3Δ,ctf3MT1*^, BJ2168^*CEN3,ctf3Δ,ctf3-MT2*^, BJ2168^*CEN3,chl4Δ,CHL4*^, BJ2168^C*EN3,chl4,chl4-MT1*^, BJ2168^*CEN3,chl4Δ,chl4-MT2*^ strains. The empty pYes2 plasmid was transformed into the BJ2168^*CEN3,cbf1Δ*^, BJ2168^*CEN3,ctf3Δ*^ and BJ2168^*CEN3,chl4Δ*^ strains as a control. Transformed yeast strains were selected on Sc-TRP-URA plates.

Single colonies of the above BJ2168 strains were cultured in Sc-URA (non-selective for mini-chromosome) for 16 h. The cultures were diluted and plated onto Sc-URA plates and incubated for 3 to 6 days at 30 °C to obtain single colonies. These colonies were restoked onto Sc-TRP-URA (yeast synthetic medium drop out tryptophan and uracil) plates, incubated for 3 to 6 days at 30 °C. Selected colonies were counted to determine the percentage of mini-chromosome retained.

#### Benomyl sensitivity assay

The method was based on published studies (Hyland et al., 1999). Freshly-grown single colonies on Sc-URA plates were suspended in water adjusted to 1x10^6^ cell/mL. The cells (in a 1/5 dilution series) were grown on YPD plus 25 μg/mL benomyl. After incubation at 25°C for 6 days, the plates were photo-recorded.

#### Immunoprecipitation and Western blotting for detecting the expression of CBF1, CENP-N and CENP-I and their respective mutants

The yeast strains were cultured in synthetic complete dropout URA and TRP media (empty pYes2-URA3 vector control), and collected at an OD^600^ of approximately 0.8. Pelleted cells were lysed in buffer (50 mM Tris.HCl pH 8.0, 300 mM NaCl, 1 mM EDTA and 1 mM DTT), and the cleared lysate was loaded onto a 1-mL Streptactin column. Fractions were eluted with 5 mM desthiobiotin and analyzed by SDS–PAGE. Western blotting was performed with a Strep-tag antibody (MCA2489P, Bio-Rad) that detected the C-terminal double StrepII-tag on CBF1, CENP-N and CENP-I. Total protein was analyzed by Coomassie blue staining for loading controls (normalized loading).

#### Cryo-EM grid preparation

For all complexes 0.05% (w/v) β-OG (n-octyl-β-D-glucopyranoside) was added to the sample immediately before plunge freezing. 3 μl of sample was applied to r2/2 Quantifoil mesh 300 grids and after 20 s of incubation, excess sample was blotted away and grids were plunge frozen in liquid ethane (blot force -10, blot time 2 s, 4°C, 100% humidity, Vitrobot markIV (ThermoFisherScientific)). The grids were screened on a 200 kV Glacios (ThermoFischerScientific) and movies were recorded on a 300 kV Titan Krios (ThermoFisherScientific) with a Falcon IV (ThermoFisherScientific) or K3 (Gatan) direct electron detector (eBIC and MRC-LMB). Data collection parameters and metrics are listed in **Table S2**.

#### Cryo-EM analysis, model building and refinement

For the CCAN-containing complexes all processing steps were carried out in Relion4.0 (Kimanius et al., 2021). Motion correction was carried out with Relion4.0, CTF estimation with CTFFIND4 (Rohou and Grigorieff, 2015). Particles were picked with Topaz (Bepler et al., 2019). After extensive 2D classification (**Figure S2**) and 3D classification 43,467 particles were used for 3D refinement of CBF1:CCAN:*153C0N3*-DNA, 100,311 particles for 3D Refinement of CBF1:CCAN:*153C0N3*-CENP-A^Nuc^:scFv and 108,672 particles for 3D refinement of CBF1:CCAN:*153C0N3*-CENP-A^Nuc^:CBF3^Core^:scFv (**Table S2**).

For the CBF1:CCAN:*153C0N3*-DNA dataset, masked 3D classification revealed a subset of 43,467 particles with well resolved density for CENP-HIK^Head^-TW (**Figure S3A**), which resulted in 3.4 Å resolution reconstruction after 3D refinement, Bayesian polishing and per-particle CTF refinement.

For the CBF1:CCAN^ΔC^:*153C0N3*-CENP-A^Nuc^:scFv dataset, consensus refinements after Bayesian polishing and per-particle CTF refinement resulted in well resolved density for CBF1:CCAN but diffuse density for the CENP-QU^Foot^ and CENP-A^Nuc^, due to conformational heterogeneity. To improve the reconstructions of our conformationally heterogeneous particle sets, we applied a variational auto-encoder that is similar to the Gaussian mixture approach proposed by (Chen and Ludtke, 2021) where conformational variability in the data is mapped to a small latent space. For a given latent coordinate, which describes the conformation of an individual particle in the data set, the decoder predicts a 3D deformation that acts on a collection of Gaussian-shaped pseudo-atoms that approximates the reconstructed density. Unique to our approach, once the 3D deformations were estimated for the entire data set, we trained a second neural network that approximates the inverse of those transformations. We then use a real-space weighted-back projection algorithm, where the original particles are back-projected along lines deformed by the inverse transformations, to obtain an improved reconstruction (details to be published elsewhere SHWS and JS) (**Figure S3B**).

For the CBF1:CCAN:*153C0N3*-CENP-A^Nuc^:CBF3^Core^:scFv data set, consensus refinements were limited to 5.6 Å resolution, locally ranging from 4.2 Å to 15 Å, due to conformational heterogeneity (**Figure S3C**). Multibody refinement with four rigid bodies was set up to increase the resolution (body 1: CCANT^op^, body 2: CCAN^Non-top^-^ΔCENP-I(Body)^, body 3: CBF3^Core^+CENP-I^Body^, body 4: CENP-A^Nuc^. All bodies refined to 3.7-3.8 Å resolution (**Figure S3C**) with clear sidechain density for most regions within each body.

For the CENP-OPQU+:CENP-A^N^ complex, micrograph movie frames were aligned with MotionCor2 and CTF estimation was performed by CTFFIND, as integrated into RELION 4.0 (Kimanius et al., 2021). Particle picking was performed using a general model in Topaz (Bepler et al., 2019). Extracted particles were initially subjected to 2D classification in cryoSPARC v3.4. *Ab initio* maps were then refined using homogeneous refinement and the resulting map was further refined using non-uniform refinement. Particles that generated the best-resolved volume were used for training a new Topaz model to improve particle picking. Newly picked particles were used as input in two rounds of heterogeneous refinement against one true map obtained from non-uniform refinement and five noisy, decoy maps, and subsequent 2D classification. The final consensus map at 3.4 Å resolution was generated through non-uniform refinement, and a small amount of anisotropy was observed.

A CBF1 monomer was modelled with Alphafold2 (Jumper et al., 2021) and docked into the cryo-EM map as a homo-dimeric bHLH with Coot (Emsley et al., 2010). Existing CCAN:CENP-A^Nuc^ (PDB ID: 6QLD) (Yan et al., 2019) and CBF3^Core^ (PDB ID: 6GYP) (Yan et al., 2018) structures were docked into the respective cryo-EM maps and adapted to fit the density with Coot. All structures were refined manually in Coot and with Phenix (Afonine et al., 2012) (**Table S2**). Figures were generated using ChimeraX (Goddard et al., 2018).

For the CENP-OPQU+:CENP-A^N^ complex, CENP-OPQU+ from the previously determined apo-CCAN structure (PDB ID 6QLF) (Yan et al., 2019) was rigid-body fitted into the CENP-OPQU+ map using Chimera. CENP-OPQU+ was then manually modified using Coot, repositioning the Nkp1-Nkp2 domain and removing flexible loops not visible in the cryo-EM density maps. The final model was refined in Phenix using default settings and model restraints from the apo-CCAN structure (PDB ID 6QLF) (Yan et al., 2019).

#### AlphaFold2 predictions

AlphaFold-Multimer (Evans et al., 2022; Jumper et al., 2021) was run to predict models for structures of the CENP-A^N^:CENP-QU and CENP-A^N^:CENP-QU-Nkp1-Nkp2 complexes. Full length CENP-Q, CENP-U, Nkp1, Nkp2, and residues of 1-120 of CENP-A were used in the prediction.

