## Supplementary Data for "Cryo-EM structure of the complete inner kinetochore of the budding yeast point centromere"

#### Supplementary Data Figure Legends

**Figure S1. Reconstituted inner kinetochore complexes.** (**A**) Alignment of DNA sequences used to reconstitute native 153 bp *CEN3*-CENP-$A^{Nuc}$ (*153CEN3*) and the near-native 153 bp *C0N3*-CENP-$A^{Nuc}$ (*153C0N3*) used in this study. The Widom 601 sequence (Lowary and Widom, 1997) is also shown. Alignment figure generated using Jalview (Waterhouse et al., 2009). (**B**) Coomassie blue-stained SDS PAGE of the *S. cerevisiae* holo-inner kinetochore complex without stabilizing scFv and with CENP-C (CBF1:CCAN:*153C0N3*-CENP-$A^{Nuc}$:$CBF3^{Core}$). (**C**) Corresponding size-exclusion chromatogram profile for sample in (B) (Agilent 1000Å column) (**D**) SEC-MALS profile of the *S. cerevisiae* holo-inner kinetochore complex. The species with mass of 1.6 MDa is consistent with a complex having a stoichiometry of 1xCBF1 dimer:2xCCAN:1x*153C0N3*-CENP-$A^{Nuc}$:1x$CBF3^{Core}$ (calculated molecular mass of 1.65 MDa). Dissociation of $CBF3^{Core}$ (240 kDa) would be consistent with a proportion of molecules having a mass of 1.3 MDa. (**E**) Coomassie blue-stained SDS PAGE of CBF1:$CCAN^{\Delta C}$:*153C0N3*-CENP-$A^{Nuc}$:scFv. (**F**) Corresponding size-exclusion chromatogram profile for sample in (E) (Agilent 1000Å column). **(G)** Coomassie blue-stained SDS PAGE of CBF1:$CCAN^{\Delta C}$:*153C0N3*-CENP-$A^{Nuc}$:$CBF3^{Core}$:scFv (inner kinetochore ($IK^{C0N3}$)). (**H**) Coomassie blue-stained SDS PAGE of CBF1:$CCAN^{\Delta C}$:*153CEN3*-CENP-$A^{Nuc}$:$CBF3^{Core}$:scFv ($IK^{CEN3}$)). (**I**) Comparative size-exclusion chromatogram profiles for samples in (G) and (H) (Agilent 1000Å column).

**Figure S2. Micrographs and 2D class averages of complexes determined in this study.** Representative micrographs and 2D class averages of (**A**) CBF1:CCAN:*153C0N3*-CENP-$A^{Nuc}$:$CBF3^{Core}$. 2D classes calculated for CBF1:CCAN:*153C0N3*-DNA. (**B**) CBF1:$CCAN^{\Delta C}$:*153C0N3*-CENP-$A^{Nuc}$:scFv and (**C**) CBF1:$CCAN^{\Delta C}$:*153C0N3*-CENP-$A^{Nuc}$ :$CBF3^{Core}$:scFv (inner kinetochore ($IK^{C0N3}$)). (**D**) In the inner kinetochore complex ($IK^{C0N3}$) a single scFv is docked on a free face of CENP-$A^{Nuc}$, no contacts between scFv and any CCAN modules were observed in the cryo-EM density or model (inset i).

**Figure S3. Local resolution maps and FSC curves of complexes determined in this study.** Local resolution estimates and FSC curves of the three reconstructions presented in this paper. (**A**) A consensus refinement of 311,873 CBF1:CCAN:*153C0N3*-DNA particles resulted in a 3.4 Å resolution reconstruction with diffuse density for CENP-$HIK^{Head}$-TW. Extensive masked 3D classification and subsequent refinement resulted in a 3.4 Å reconstruction, from 43,467 particles where side-chains were resolved for CENP-$HIK^{Head}$-TW. (**B**) A consensus refinement of 100,311 CBF1:CCAN:*153C0N3*-CENP-$A^{Nuc}$:scFv particles resulted in a 3.7 Å resolution reconstruction with well-defined density for CCAN while the CENP-$A^{Nuc}$ and CENP-$QU^{Foot}$ modules were resolved at significantly lower resolution. After deformation refinement with an in-house algorithm (Methods) the local resolutions of CENP-$A^{Nuc}$ and CENP-$QU^{Foot}$ increased significantly, as well as the global resolution, now at 3.4 Å. At this improved resolution the model major and minor grooves of the *C0N3* DNA duplex were resolved, allowing for accurate mapping of the dyad. (**C**) 108,672 particles contributed to a 5.6 Å resolution consensus reconstruction of the inner kinetochore complex - CBF1:$CCAN^{\Delta C}$:*153C0N3*-CENP-$A^{Nuc}$:$CBF3^{Core}$:scFv ($IK^{C0N3}$). (**D**) Multibody refinement with four bodies ($CCAN^{Top}$, $CCAN^{Non\text{-}top\text{-}\Delta CENP\text{-}I\text{-}Body}$, $CBF3^{Core}$+CENP-$I^{Body}$ and CENP-$A^{Nuc}$) increased the resolution locally to 3.7-3.8 Å for each body.

**Figure S4. (A) CBF1 interacts with the back-face of $CCAN^{Top}$.** Details of how CBF1 interacts with CENP-P (inset i), CENP-Q (inset ii), CENP-N (inset iii) and CENP-L (inset iv). (**B**) Western blot showing that wildtype and mutant CBF1, CENP-N and CENP-I are expressed at equivalent levels in the BJ2168$^{CEN3}$ strain (left) and loading control (right; Coomassie-blue-stained gel shows dynein).

**Figure S5. Structures of CBF3$^{Core}$ and model of inner kinetochore-CENP-A$^{Nuc}$ complex with CENP-C and Ndc10$^{DBD}$.** (**A**) and (**B**) Comparison of the binding mode of CBF3$^{Core}$ to CENP-A$^{Nuc}$ in isolation (Guan et al., 2021) (A), and in the context of the inner kinetochore (this work) (B). CBF3$^{Core}$ rotates around the flexible hinge linking it to the Gal4 domain when in the context of the inner kinetochore. (**C**) A model showing the inner kinetochore-CENP-A$^{Nuc}$ complex represented by a surface and the CENP-C dimer (residues 222 to C-term) and Ndc10$^{DBD}$ (DNA-binding domain of Ndc10, residues 1-540) in cartoon. Ndc10$^{DBD}$ was modelled based on the cryo-EM structure of CBF3$^{Holo}$ (Yan et al., 2018) (PDB 6GYP). The model of CENP-C is based on the crystal structure of *S. cerevisiae* CENP-C$^{Cupin}$ (Cohen et al., 2008) and the cryo-EM structure of the CENP-C motif in complex with CENP-A$^{Nuc}$ (PDB 7ON1), combined with the AlphaFold2 prediction of *S. cerevisiae* CENP-C (Tunyasuvunakool et al., 2021). Regions of CENP-C linking its MIND-binding motif with residue 260 are predicted to be disordered. Residues 1 to 221 are not shown. In the model, residues 327-340 are predicted to form a basic α-helix (CENP-C$^{\alpha2}$) that inserts into the DNA major groove of CDEII. This α-helix is within the defined DNA-binding domain (residues 256-356), and is consistent with data that CENP-C interacts with a dyad-adjacent site within CDEII (Xiao et al., 2017). The CENP-C motif (residues 282-305), is shown in yellow. Residues 222-240 of CENP-C are in close proximity to the CENP-Q$^{192-220}$ α-helix shown in blue. These regions of CENP-C and CENP-Q were shown to interact by CLMS (Fischbock-Halwachs et al., 2019), and deletion of either region abolished phospho-CENP-C association with CENP-QU (Hagemann et al., 2022). The CENP-LN and CENP-HIKM-binding motifs of human CENP-C (Klare et al., 2015; Pentakota et al., 2017; Pesenti et al., 2022; Tian et al., 2018; Yatskevich et al., 2022) are not conserved in *S. cerevisiae* CENP-C/Mif2p. Likewise, the CENP-C-binding sites on human CENP-LN and CENP-HIKM (Pesenti et al., 2022; Tian et al., 2018; Yatskevich et al., 2022) are not present in *S. cerevisiae* CCAN. (**D**) Close-up of the CEP3$^{Gal4}$ density in the cryo-EM map at lower threshold. (**E**) and (**F**). Comparison of *CEN3*-CENP-A$^{Nuc}$ (Guan et al., 2021) (E) with *153C0N3*-CENP-A$^{Nuc}$ as part of the inner kinetochore from this study (F). CDEI (disordered in the *CEN3*-CENP-A$^{Nuc}$ structure) is exposed in the inner kinetochore due to unwrapping of the 5'-end. Likewise, the 3' end is slightly more unwrapped in the inner kinetochore compared to the *CEN3*-CENP-A$^{Nuc}$ and engaged by CCAN$^{Non\text{-}top}$.

**Figure S6. Gallery of 2D classes representing the structural states of CCAN species. (A-C)** 2D class averages from cryo-EM data sets for (A) CBF1:CCAN$^{\Delta C}$:*153C0N3*-CENP-A$^{Nuc}$:scFv, (B) CBF1:CCAN$^{\Delta C}$:*153C0N3*-CENP-A$^{Nuc}$:CBF3$^{Core}$:scFv (IK$^{C0N3}$) and (C) CBF1:CCAN$^{\Delta C}$:*153CEN3*-CENP-A$^{Nuc}$:CBF3$^{Core}$:scFv (IK$^{CEN3}$). Shown are four sets of 2D class averages represented in the three data sets: (i) Dimeric CCAN (DNA, no CENP-A$^{Nuc}$), similar to the apo-dimeric CCAN reported by (Hinshaw and Harrison, 2019), (ii) monomeric CCAN:CENP-A$^{Nuc}$, (iii) pseudo-symmetric and (iv) asymmetric di-CCAN configurations assembled on CENP-A$^{Nuc}$. (**A**) In the CBF1:CCAN$^{\Delta C}$:*153C0N3*-CENP-A$^{Nuc}$:scFv dataset, only ~0.5% of the particles correspond to a dimeric CCAN:DNA species, devoid of CENP-A$^{Nuc}$ (i), while ~40% of the particles correspond to monomeric CBF1:CCAN:*153C0N3*-CENP-A$^{Nuc}$ with a CCAN$^{Top}$ configuration (ii). ~10% of particles correspond to pseudo-symmetric di-CCAN$^{Non\text{-}top}$ assembled on CENP-A$^{Nuc}$ (iii). Only ~1% of particles correspond to an asymmetric di-CCAN species assembled on CENP-A$^{Nuc}$, with one CCAN$^{Top}$ and one CCAN$^{Non\text{-}top}$. (**B**) In the CBF1:CCAN$^{\Delta C}$:*153C0N3*-CENP-A$^{Nuc}$:CBF3$^{Core}$:scFv (IK$^{C0N3}$) sample, no dimeric CCAN:DNA species were observed (i), and significantly less monomeric CBF1:CCAN$^{\Delta C}$:*153C0N3*-CENP-A$^{Nuc}$ (ii) and pseudo-symmetric di-CCAN$^{Non\text{-}top}$ assembled on

CENP-A$^{Nuc}$ (iii). Instead, the equilibrium is shifted towards the asymmetric di-CCAN species assembled on CENP-A$^{Nuc}$ of the inner kinetochore (~14% of particles) (iv). (**C**) In the CBF1:CCAN$^{\Delta C}$:*153CEN3*-CENP-A$^{Nuc}$:CBF3$^{Core}$:scFv sample, ~13% of particles correspond to a dimeric CCAN:DNA species, devoid of CENP-A$^{Nuc}$ (i), while only 3.2% of particles correspond to the pseudo-symmetric di-CCAN$^{Non\text{-}top}$ species assembled on CENP-A$^{Nuc}$ (iii), and 1.4% correspond to asymmetric di-CCAN species as observed in the inner kinetochore (iv). No monomeric CCAN:CENP-A$^{Nuc}$ species were observed (ii). These numbers are indicative of poor *CEN3*-CENP-A$^{Nuc}$ stability in solution and/or on cryo-EM grids. The CCAN modules in the pseudo-symmetric di-CCAN species adopt a range of conformational states with respect to CENP-A$^{Nuc}$ from more closed to more open, as is apparent from the 2D class averages in column iii. Particle fractions are calculated as a percentage of total extracted particles per dataset. The remaining extracted particles were denatured and/or false positives. (**D**) A 10 Å-resolution3D reconstruction of a symmetric di-CCAN:*CEN3*-CENP-A$^{Nuc}$:CBF1 assembly, where both CCAN protomers bind the *CEN3* DNA non-topologically in the absence of CBF3$^{Core}$. scFv binds to one face of CENP-A$^{Nuc}$. The identical reconstruction was obtained from the CBF1:CCAN$^{\Delta C}$*:153C0N3*-CENP-A$^{Nuc}$:CBF3$^{Core}$:scFv (IK$^{C0N3}$) cryo-EM data set.

**Figure S7. The N-terminus of CENP-A (CENP-A$^{N}$) interacts with CENP-QU.** (**A**) Sequences of CENP-A$^{END}$ peptides modelled on the N-terminus of CENP-A used in this study. Alignment figure generated using Jalview (Waterhouse et al., 2009). (**B**) Quantification of isothermal titration calorimetry data shown in panels C-E. (**C-E**) Isothermal titration calorimetry measurements of the binding of CENP-A$^{END}$ peptides (CENP-A$^{END\text{-}1}$, CENP-A$^{END\text{-}2}$, CENP-A$^{END\text{-}3}$, respectively) to CENP-QU. Upper panel, raw data of the titration of CENP-A$^{END}$ into CENP-QU. Lower panel, integrated heats of injections, corrected for the heat of dilution, with the solid line corresponding to the best fit of the data using the MicroCal software. (**F**) CENP-QU and CENP-A$^{N}$ form a stable complex, as judged by SEC. Upper panel: SEC chromatogram (Superdex S75 column), lower panel: corresponding Coomassie-blue stained PAGE. (**G**) Mutating residues in CENP-QU (D191K$^{CENP\text{-}U}$, D194K$^{CENP\text{-}U}$ and E235K$^{CENP}$) predicted to interact with R37, R46 and K49 of CENP-A$^{N}$ disrupts the CENP-QU:CENP-A$^{N}$ complex. Upper panel: SEC chromatogram (Superdex S75 column), lower panel: corresponding Coomassie-blue stained PAGE.

**Figure S8. Confidence metrics for the CENP-A$^{N}$:CENP-QU AlphaFold2 prediction**. (**A**) Predicted local distance difference test (lDDT) values (Mariani et al., 2013) for the AlphaFold2 model of CENP-A$^{N}$:CENP-QU. A higher score indicates higher confidence in the prediction. **(B)** The predicted alignment error (PAE) heat map for the CENP-A$^{N}$:CENP-QU AlphaFold2 prediction. The PAE heat map shows the predicted error (in angstroms) between all pairs of residues, with blue indicating lower error and red indicating higher error. The PAE plot suggests high confidence for the predicted interactions between CENP-QU and CENP-A$^{END}$. (**C**) The pLDDT values mapped onto the predicted model of CENP-A$^{N}$:CENP-QU. Residues are colored in a scale of high (red) to low (blue) pLDDT values. The model of CENP-QU bound to CENP-A$^{END}$ was predicted with high confidence (pLDDT > 90), whereas the residues 65-97 of CENP-A were predicted with relatively low confidence (pLDDT < 60). (**D**) Top panel: Coomassie blue-stained PAGE of the CENP-OPQU+:CENP-A$^{N}$ complex, below: corresponding SEC chromatogram (Agilent 1000Å column). (**E**) Left: cryo-EM map of the CENP-OPQU+:CENP-A$^{N}$ complex color-coded according to local resolution. Right: FSC curves of the CENP-OPQU+:CENP-A$^{N}$ complex (green (unmasked), blue (masked)).

**Figure S9. CENP-A$^{N}$-binding to CCAN is auto-inhibited and supernumerary CENP-QU interacts with the inner kinetochore:CENP-A$^{Nuc}$ complex through CENP-A$^{N}$.** (**A**) CENP-A$^{N}$ (residues 1-82) does not bind to CCAN. Upper panel: Coomassie-blue stained PAGE, lower panel: corresponding SEC chromatogram (Agilent 1000Å column). (**B**) The binding of CENP-A$^{N}$ to CENP-OPQU+ is displaced by CENP-LN. In the presence of CENP-A$^{N}$ and CENP-LN, CENP-OPQU+ forms either CENP-OPQU+:CENP-A$^{N}$ or CENP-OPQU+:CENP-LN complexes, showing that CENP-A$^{N}$ and CENP-LN binding to CENP-OPQU+ is mutually exclusive. Upper panel: Coomassie-blue stained PAGE, lower panel: corresponding SEC chromatogram (Superose 6 column). (**C**) Coomassie-blue stained PAGE of SEC fractions (Agilent 1000Å column) showing binding of supernumerary CENP-QU to the inner kinetochore:CENP-A$^{Nuc}$ complex. The addition CENP-ΔQU complex (CENP-ΔQ, CENP-ΔU) consisted of residues 30-255 (CENP-ΔQ) and 1-294 (CENP-ΔU) to distinguish from CENP-QU assembled into CCAN. (**D**) Binding of supernumerary CENP-QU is abolished when CENP-A$^{\Delta N}$ (deletion of CENP-A residues 1-129) was used to reconstitute CENP-A$^{Nuc}$ (CENP-A$^{Nuc-\Delta N}$). Coomassie-blue stained PAGE of SEC fractions (Agilent 1000Å column).

**Table S1. Kinetochore subunit and sub-complex nomenclature for *S. cerevisiae* and human.**

**Table S2. Cryo-EM data collection, refinement and validation statistics.**

**Video S1. Structure of the S. cerevisiae inner kinetochore on a CENP-A nucleosome.** The video shows how the component sub-complexes of the inner kinetochore are arranged on a central CENP-A nucleosome to build the entire inner kinetochore-CENP-A$^{Nuc}$ complex. The extensive unwrapping of CENP-A$^{Nuc}$ DNA ends creates the binding sites for two CCAN protomers. The organization of CCAN$^{Top}$ and CCAN$^{Non-top}$ is assisted by two DNA-specific binding complexes, CBF1 and CBF3, that engage, respectively, the conserved CDEI and CDEIII sequence elements conserved in point centromeres.

**Video S2. Model for CENP-A essential N-terminal domain (CENP-A$^{END}$) interaction with CENP-QU.** The interaction of CENP-A$^{END}$ with CENP-QU requires that Nkp1-Nkp2 undergo a conformational change to expose the CENP-A$^{END}$-binding site on CENP-QU.

#### Table S1 Nomenclature and organization of kinetochore subunits and sub-complexes

| Subunit | *S.c.* name | Length | Mol mass kDa | Sub-complex | Level2 | Level3 |
|---|---|---|---|---|---|---|
| ***153C0N3*-CENP-A nucleosome** | | | | **CENP-A$^{Nuc}$** | **Inner kinetochore** | **Holo-kinetochore** |
| **CENP-A** | Cse4 | 229 | 26.8 | | | |
| **H2A** | HTA2 | 132 | 14.0 | | | |
| **H2B** | HTB1 | 131 | 14.2 | | | |
| **H4** | HHF1 | 103 | 11.4 | | | |
| **1*53C0N3* DNA** | | 153 bp | 94.9 | | | |
| **CENP-C** | | | | **CENP-C** | | |
| **CENP-C** | Mif2 | 549 | 62.5 | | | |
| **CENP-HIK-TW complex** | | | | **CCAN** | | |
| **CENP-H** | Mcm16 | 181 | 21.1 | | | |
| **CENP-I** | Ctf3 | 733 | 84.3 | | | |
| **CENP-K** | Mcm22 | 239 | 27.6 | | | |
| **CENP-T** | Cnn1 | 361 | 41.3 | | | |
| **CENP-W** | Wip1 | 89 | 10.2 | | | |
| **CENP-LN complex** | | | | | | |
| **CENP-L** | Iml3 | 245 | 28.1 | | | |
| **CENP-N** | Chl4 | 458 | 52.7 | | | |
| **CENP-OPQU+/COMA+ complex** | | | | | | |
| **CENP-O** | Mcm21 | 368 | 43.0 | | | |
| **CENP-P** | Ctf19 | 369 | 42.8 | | | |
| **CENP-Q** | Okp1 | 406 | 47.3 | | | |
| **CENP-U** | Ame1 | 324 | 37.5 | | | |
| **Nkp1** | Nkp1 | 238 | 27.0 | | | |
| **Nkp2** | Nkp2 | 153 | 17.9 | | | |
| **CBF3$^{Core}$ complex** | | | | **CBF3** | | |
| **Cep3** | Cep3 | 608 | 71.4 | | | |
| **Ctf13** | Ctf13 | 478 | 56.3 | | | |
| **Skp1** | Skp1 | 194 | 22.3 | | | |
| **Ndc10** | Ndc10 | 956 | 111.9 | | | |
| **CBF1 complex** | | | | **CBF1** | | |
| **Cbf1** | Cbf1 | 351 | 39.4 | | | |
| **Ndc80 complex** | | | | **NDC80c** | **KMN network of outer kinetochore** | |
| **Ndc80** | Ndc80 | 691 | 80.5 | | | |
| **Nuf2** | Nuf2 | 451 | 53.0 | | | |
| **Spc24** | Spc24 | 213 | 24.6 | | | |
| **Spc25** | Spc25 | 221 | 25.2 | | | |
| **MIND/MTW1 complex** | | | | **MIND** | | |
| **Dsn1** | Dsn1 | 576 | 65.7 | | | |
| **Nnf1** | Nnf1 | 201 | 23.6 | | | |
| **Mis12** | Mtw1 | 289 | 33.2 | | | |
| **Nsl1** | Nsl1 | 216 | 25.4 | | | |
| **KNL1 complex** | | | | **KNL1c** | | |
| **Knl1** | Spc105 | 917 | 104.8 | | | |
| **Zwint1** | Kre28 | 385 | 44.7 | | | |
| **scFv** | - | 265 | 29.0 | NA | NA | |

#### Table S2 Cryo-EM data collection, refinement and validation statistics

| | CBF1: CCAN: *153C0N3* (EMDB-xxxx) (PDB xxxx) | CBF1: CCAN: *153C0N3*-CENP-A$^{Nuc}$ (EMDB-xxxx) (PDB xxxx) | CBF1: CCAN: *153C0N3*-CENP-A$^{Nuc}$: CBF3$^{Core}$ (IK$^{C0N3}$) (EMDB-xxxx) (PDB xxxx) | CENP-OPQU+: CENP-A$^{N}$ (EMDB-xxxx) (PDB xxxx) |
|---|---|---|---|---|
| **Data collection and processing** | | | | |
| Magnification | 105,000 | 105,000 | 105,000 | 105,000 |
| Voltage (kV) | 300 | 300 | 300 | 300 |
| Electron exposure (e–/Å$^2$) | 40 | 40 | 40 | 40 |
| Detector | Falcon4 | K3 | Falcon4 | K3 |
| Defocus range (μm) | 1.2-2.6 | 1.2-2.6 | 1.2-2.6 | 0.5-2.0 |
| Pixel size (Å) | 1.17 | 0.853 | 1.08 | 0.826 |
| Symmetry imposed | C1 | C1 | C1 | C1 |
| Movies collected (N) | 8,804 | 10,385 | 10,196 | 12,414 |
| Initial particle (N) | 1,516,955 | 1,142,899 | 1,217,837 | 3,673,108 |
| Final particle images (N) | 43,467 | 100,311 | 108,672 | 595,147 |
| Map resolution (Å) | 3.4 | 3.4 | 3.7-3.8 | 3.4 |
| FSC threshold | 0.143 | 0.143 | 0.143 | 0.143 |
| Map resolution range (Å) | 3.4-8 | 3.4-9.6 | 3.4-6.8 | 3.4-12 |
| Map sharpening *B* factor (Å$^2$) | -85 | -110 | -110 | -70 |
| **Refinement** | | | | |
| Initial model used (PDB code) | *6QLE, Ab initio* | *6QLD* | *6QLD,6GYP* | *6QLF* |
| Model resolution (Å) | 3.4 | 3.5 | 3.7 | 7.0 |
| FSC threshold | 0.5 | 0.5 | 0.5 | 0.5 |
| Model to map CC | 0.84 | 0.76 | 0.71 | 0.6 |
| Model composition | | | | |
| Non-hydrogen atoms (N) | 28,484 | 37,995 | 78,805 | 67,63 |
| Protein residues (N) | 3,384 | 4,108 | 8,937 | 885 |
| Nucleotides (N) | 54 | 241 | 306 | 0 |
| **Validation** | | | | |
| *B* factors (Å$^2$) | | | | |
| Protein | 96.15 | 138.11 | 109.89 | 83.44 |
| Nucleotide | 132.94 | 175.66 | 152.19 | N/A |
| R.m.s. deviations | | | | |
| Bond lengths (Å) | 0.004 | 0.004 | 0.004 | 0.003 |
| Bond angles (°) | 0.665 | 0.732 | 0.727 | 0.559 |
| Validation | | | | |
| MolProbity score | 1.89 | 1.92 | 1.75 | 1.98 |
| Clash score | 12.77 | 13.22 | 8.52 | 14.21 |
| Poor rotamers (%) | 0.10 | 0.41 | 1.00 | 0.28 |
| Ramachandran plot | | | | |
| Favoured (%) | 96.00 | 95.88 | 95.87 | 95.43 |
| Allowed (%) | 3.97 | 4.12 | 4.09 | 4.57 |
| Disallowed (%) | 0.03 | 0.00 | 0.03 | 0.00 |

### Figure S1

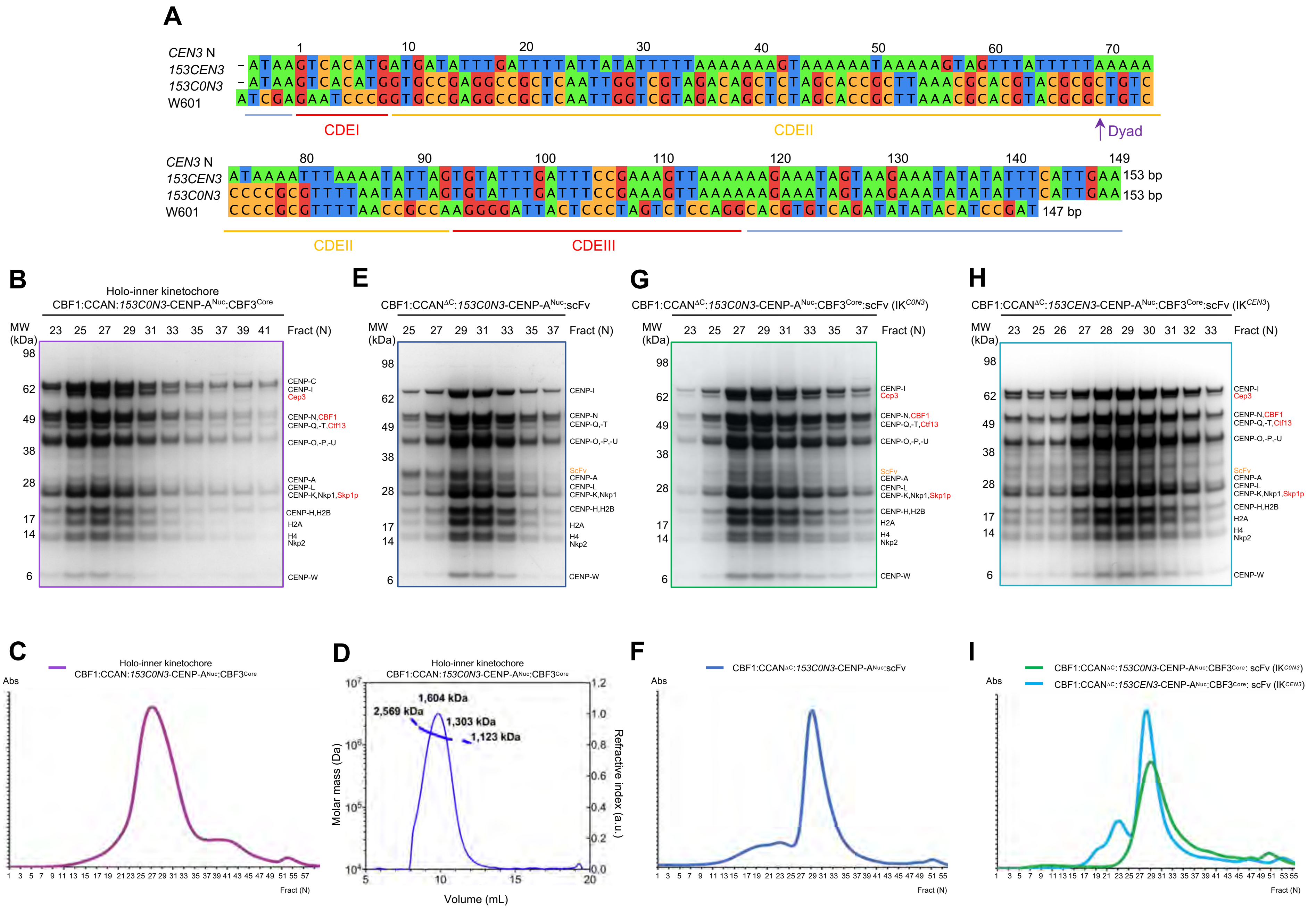

A
CEN3 N
153CEN3
153C0N3
W601
CDEI
CDEII
Dyad
CDEIII
153 bp
153 bp
147 bp
B
Holo-inner kinetochore
CBF1:CCAN:153C0N3-CENP-ANuc:CBF3Core
E
CBF1:CCANΔC:153C0N3-CENP-ANuc:scFv
G
CBF1:CCANΔC:153C0N3-CENP-ANuc:CBF3Core:scFv (IKC0N3)
H
CBF1:CCANΔC:153CEN3-CENP-ANuc:CBF3Core:scFv (IKCEN3)
MW (kDa)
Fract (N)
CENP-C
CENP-I
Cep3
CENP-N,CBF1
CENP-Q,-T,Ctf13
CENP-O,-P,-U
ScFv
CENP-A
CENP-L
CENP-K,Nkp1,Skp1p
CENP-H,H2B
H2A
H4
Nkp2
CENP-W
C
Holo-inner kinetochore
CBF1:CCAN:153C0N3-CENP-ANuc:CBF3Core
Abs
D
Holo-inner kinetochore
CBF1:CCAN:153C0N3-CENP-ANuc:CBF3Core
1,604 kDa
2,569 kDa
1,303 kDa
1,123 kDa
Molar mass (Da)
Refractive index (a.u.)
Volume (mL)
F
CBF1:CCANΔC:153C0N3-CENP-ANuc:scFv
I
CBF1:CCANΔC:153C0N3-CENP-ANuc:CBF3Core: scFv (IKC0N3)
CBF1:CCANΔC:153CEN3-CENP-ANuc:CBF3Core: scFv (IKCEN3)
Fract (N)

**Figure S2**

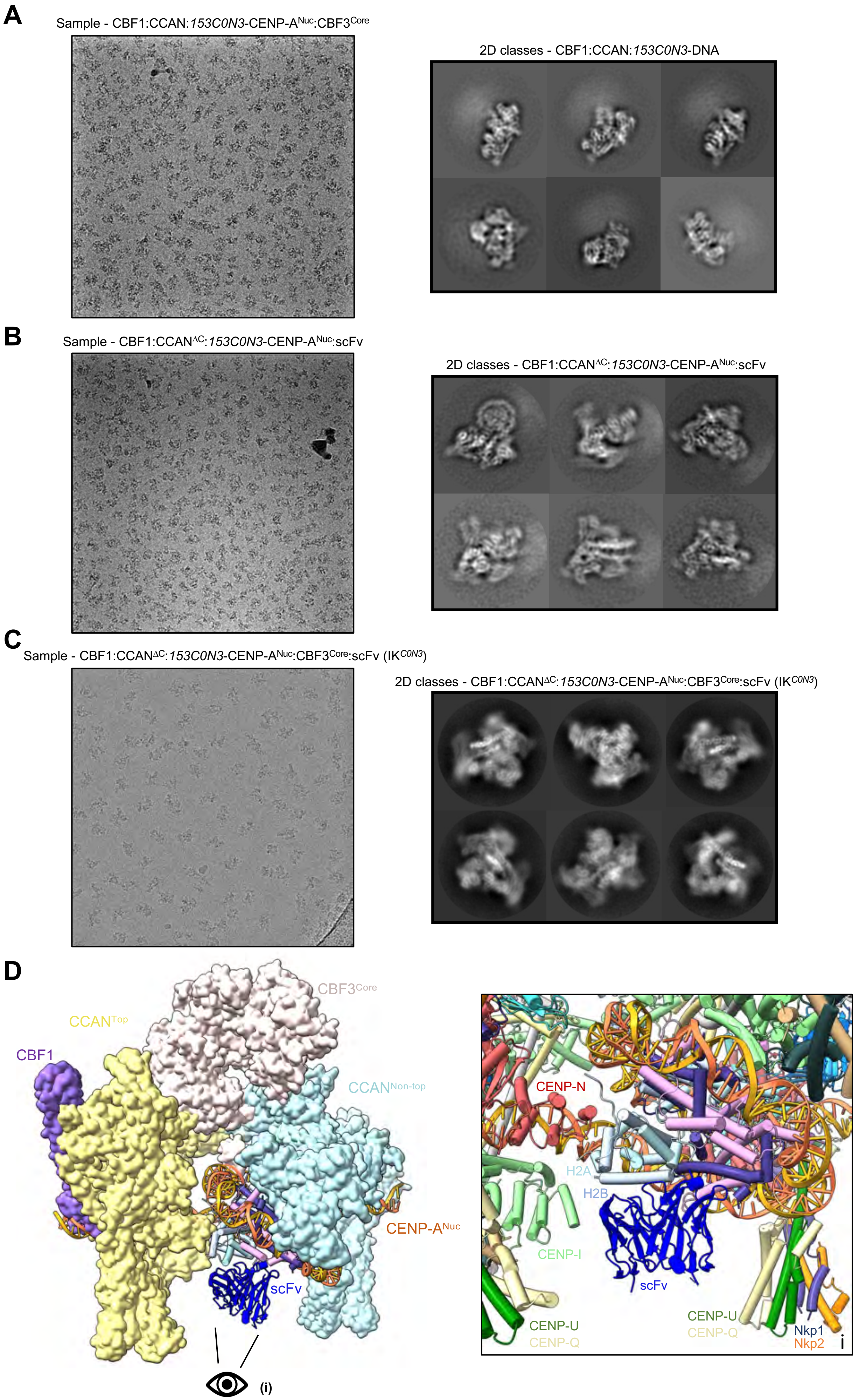
A
Sample - CBF1:CCAN:153C0N3-CENP-ANuc:CBF3Core
2D classes - CBF1:CCAN:153C0N3-DNA
B
Sample - CBF1:CCANΔC:153C0N3-CENP-ANuc:scFv
2D classes - CBF1:CCANΔC:153C0N3-CENP-ANuc:scFv
C
Sample - CBF1:CCANΔC:153C0N3-CENP-ANuc:CBF3Core:scFv (IKC0N3)
2D classes - CBF1:CCANΔC:153C0N3-CENP-ANuc:CBF3Core:scFv (IKC0N3)
D
CBF3Core
CCANTop
CBF1
CCANNon-top
CENP-ANuc
scFv
(i)
CENP-N
H2A
H2B
CENP-I
scFv
CENP-U
CENP-Q
CENP-U
CENP-Q
Nkp1
Nkp2
i

### Figure S3

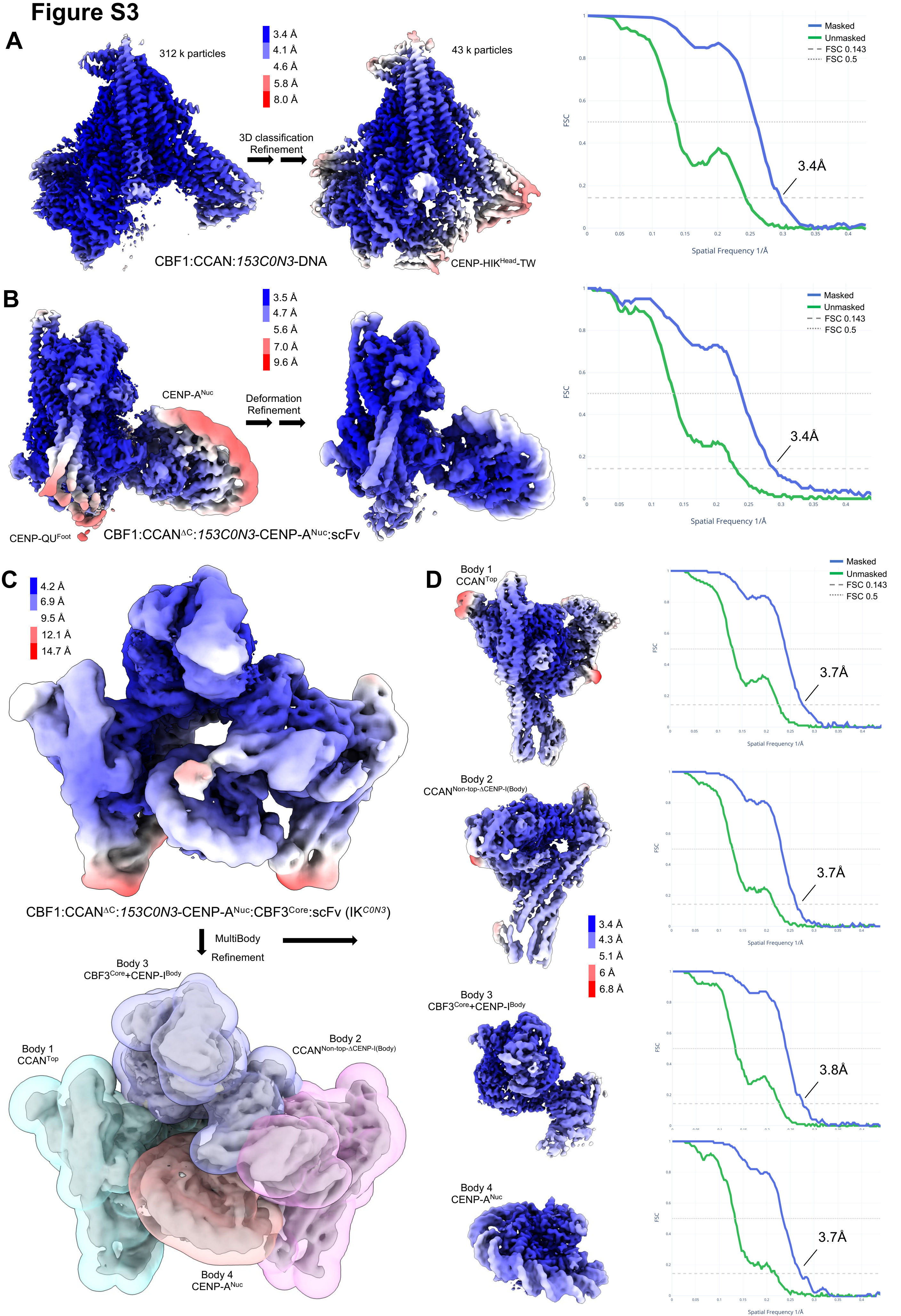
A
312 k particles
3.4 Å
4.1 Å
4.6 Å
5.8 Å
8.0 Å
3D classification
Refinement
43 k particles
CBF1:CCAN:153C0N3-DNA
CENP-HIK^Head-TW
Masked
Unmasked
FSC 0.143
FSC 0.5
FSC
Spatial Frequency 1/Å
3.4Å
B
3.5 Å
4.7 Å
5.6 Å
7.0 Å
9.6 Å
CENP-A^Nuc
Deformation
Refinement
CENP-QU^Foot
CBF1:CCAN^ΔC:153C0N3-CENP-A^Nuc:scFv
3.4Å
C
4.2 Å
6.9 Å
9.5 Å
12.1 Å
14.7 Å
CBF1:CCAN^ΔC:153C0N3-CENP-A^Nuc:CBF3^Core:scFv (IK^C0N3)
MultiBody
Refinement
Body 3
CBF3^Core+CENP-I^Body
Body 1
CCAN^Top
Body 2
CCAN^Non-top-ΔCENP-I(Body)
Body 4
CENP-A^Nuc
D
Body 1
CCAN^Top
3.7Å
Body 2
CCAN^Non-top-ΔCENP-I(Body)
3.7Å
3.4 Å
4.3 Å
5.1 Å
6 Å
6.8 Å
Body 3
CBF3^Core+CENP-I^Body
3.8Å
Body 4
CENP-A^Nuc
3.7Å

### Figure S4

**A**

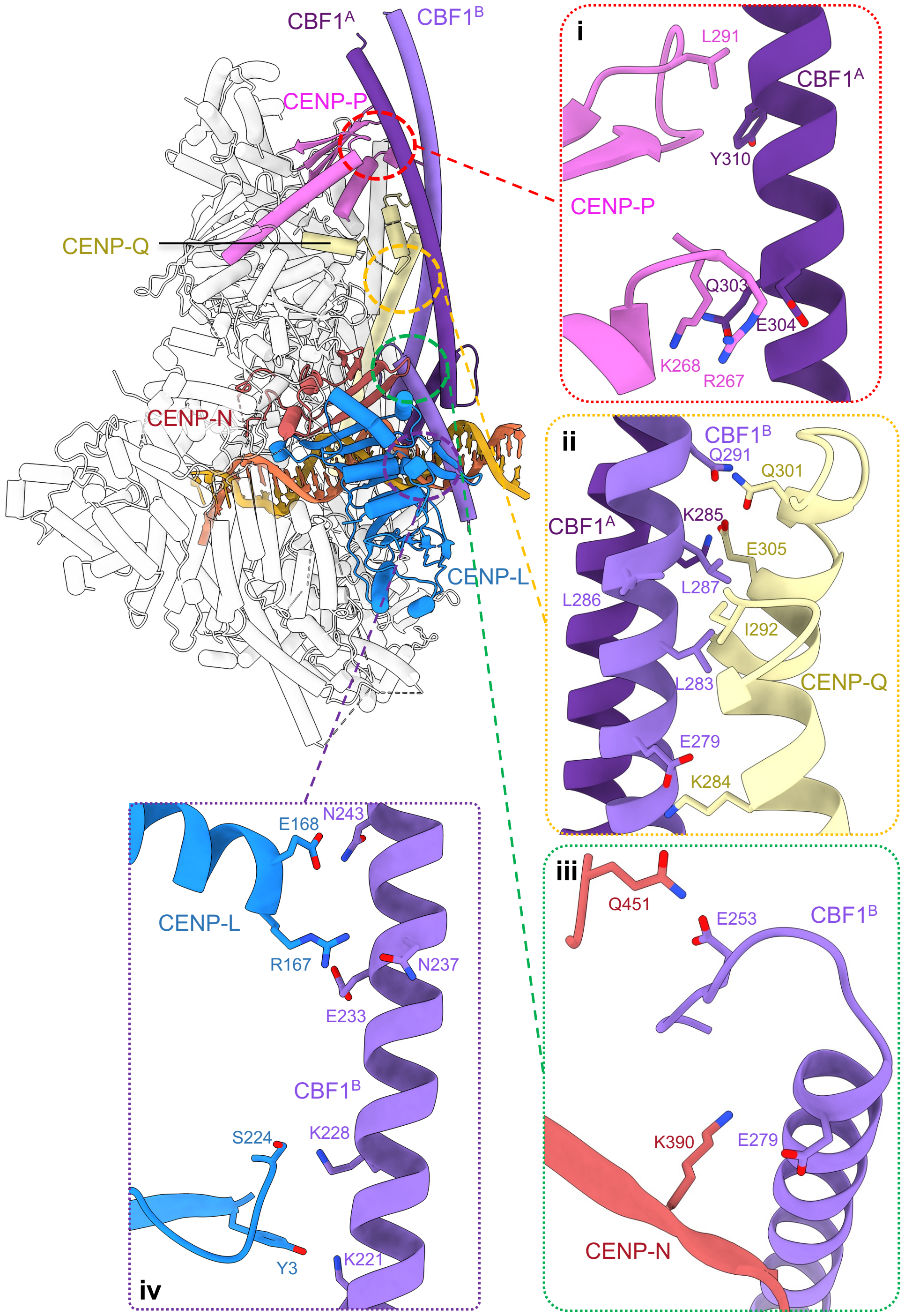

CBF1$^{A}$
CBF1$^{B}$
CENP-P
CENP-Q
CENP-N
CENP-L
i
L291
CBF1$^{A}$
Y310
CENP-P
Q303
E304
K268
R267
ii
CBF1$^{B}$
Q291
Q301
CBF1$^{A}$
K285
E305
L287
L286
I292
L283
CENP-Q
E279
K284
iii
Q451
E253
CBF1$^{B}$
K390
E279
CENP-N
iv
N243
E168
CENP-L
R167
N237
E233
CBF1$^{B}$
S224
K228
Y3
K221

**B**

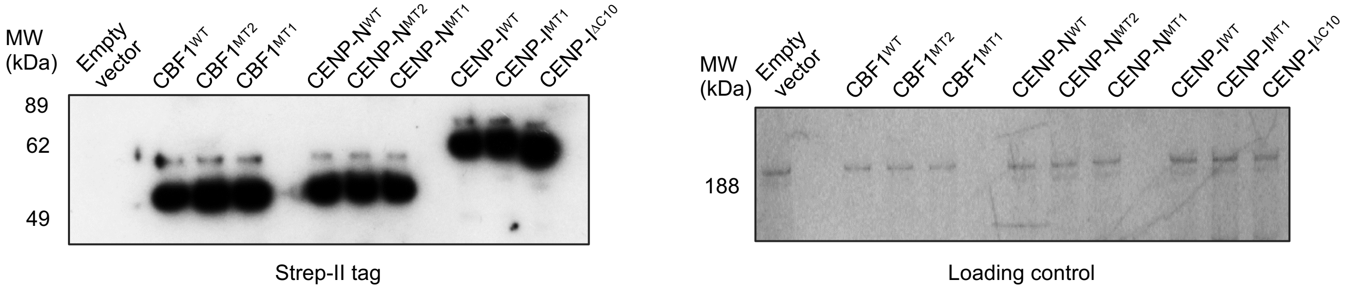

MW (kDa)
Empty vector
CBF1$^{WT}$
CBF1$^{MT2}$
CBF1$^{MT1}$
CENP-N$^{WT}$
CENP-N$^{MT2}$
CENP-N$^{MT1}$
CENP-I$^{WT}$
CENP-I$^{MT1}$
CENP-I$^{\Delta C10}$
89
62
49
Strep-II tag
188
Loading control

**Figure S5**

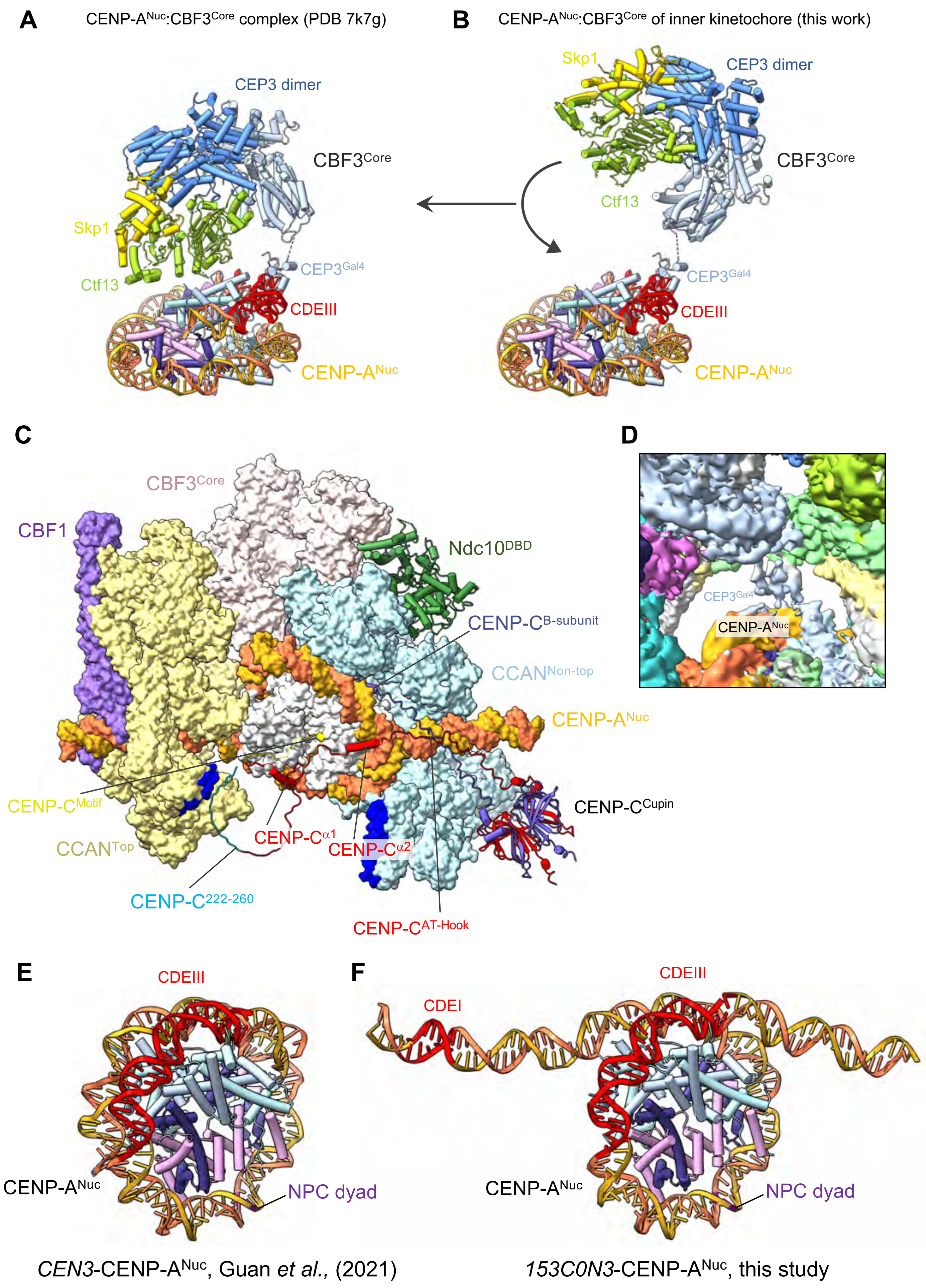
A
CENP-A$^{Nuc}$:CBF3$^{Core}$ complex (PDB 7k7g)
CEP3 dimer
CBF3$^{Core}$
Skp1
Ctf13
CEP3$^{Gal4}$
CDEIII
CENP-A$^{Nuc}$
B
CENP-A$^{Nuc}$:CBF3$^{Core}$ of inner kinetochore (this work)
Skp1
CEP3 dimer
CBF3$^{Core}$
Ctf13
CEP3$^{Gal4}$
CDEIII
CENP-A$^{Nuc}$
C
CBF3$^{Core}$
CBF1
Ndc10$^{DBD}$
CENP-C$^{B\text{-}subunit}$
CCAN$^{Non\text{-}top}$
CENP-A$^{Nuc}$
CENP-C$^{Motif}$
CCAN$^{Top}$
CENP-C$^{\alpha1}$
CENP-C$^{\alpha2}$
CENP-C$^{Cupin}$
CENP-C$^{222\text{-}260}$
CENP-C$^{AT\text{-}Hook}$
D
CEP3$^{Gal4}$
CENP-A$^{Nuc}$
E
CDEIII
CENP-A$^{Nuc}$
NPC dyad
*CEN3*-CENP-A$^{Nuc}$, Guan *et al.,* (2021)
F
CDEIII
CDEI
CENP-A$^{Nuc}$
NPC dyad
*153C0N3*-CENP-A$^{Nuc}$, this study

### Figure S6

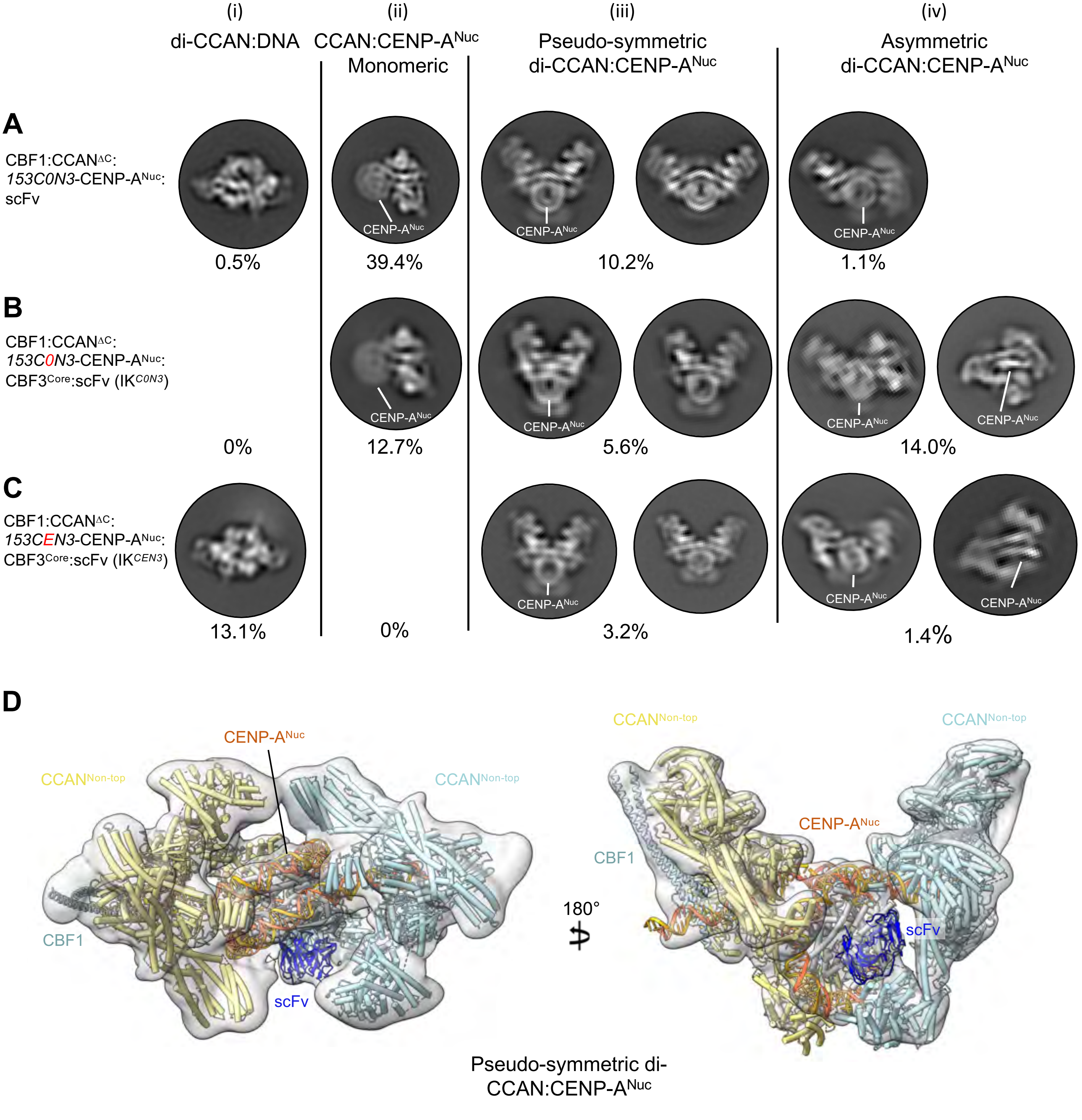

(i)
di-CCAN:DNA
(ii)
CCAN:CENP-A^Nuc
Monomeric
(iii)
Pseudo-symmetric
di-CCAN:CENP-A^Nuc
(iv)
Asymmetric
di-CCAN:CENP-A^Nuc
A
CBF1:CCAN^ΔC:
153C0N3-CENP-A^Nuc:
scFv
CENP-A^Nuc
0.5%
39.4%
10.2%
1.1%
B
CBF1:CCAN^ΔC:
153C0N3-CENP-A^Nuc:
CBF3^Core:scFv (IK^C0N3)
0%
12.7%
5.6%
14.0%
C
CBF1:CCAN^ΔC:
153CEN3-CENP-A^Nuc:
CBF3^Core:scFv (IK^CEN3)
13.1%
0%
3.2%
1.4%
D
CENP-A^Nuc
CCAN^Non-top
CBF1
scFv
180°
Pseudo-symmetric di-
CCAN:CENP-A^Nuc

### Figure S7

**A**

```
                      32   37   42   47   52   57
CENP-A END-1  - - - - - S I N D R A L S L L Q R T R A T - - - - - - - - - - - -
CENP-A END-2  A G D Q Q S I N D R A L S L L Q R T R A T K N - - - - - - - - - - -
CENP-A END-3  A G D Q Q S I N D R A L S L L Q R T R A T K N L F P R R E E R R R
```

**B**

| Peptide | N | Kd (μM) | ΔH kcal$^{-1}$ | -TΔS kcal$^{-1}$ | ΔG kcal$^{-1}$ | [protein] mM | [peptide] mM |
|---|---|---|---|---|---|---|---|
| CENP-A$^{END-1}$ | 0.99 ± 0.01 | 11.63 ± 0.19 | -11.47 ± 0.05 | 4.84 ± 0.03 | -6.62 ± 0.01 | 0.180 | 2.18 |
| CENP-A$^{END-2}$ | 0.95 ± 0.01 | 1.09 ± 0.07 | -12.75 ± 0.05 | 4.74 ± 0.10 | -8.0 ± 0.04 | 0.142 | 1.24 |
| CENP-A$^{END-3}$ | 1.09 ± 0.01 | 0.72 ± 0.03 | -13.10 ± 0.10 | 4.87 ± 0.12 | -8.25 ± 0.03 | 0.142 | 1.18 |

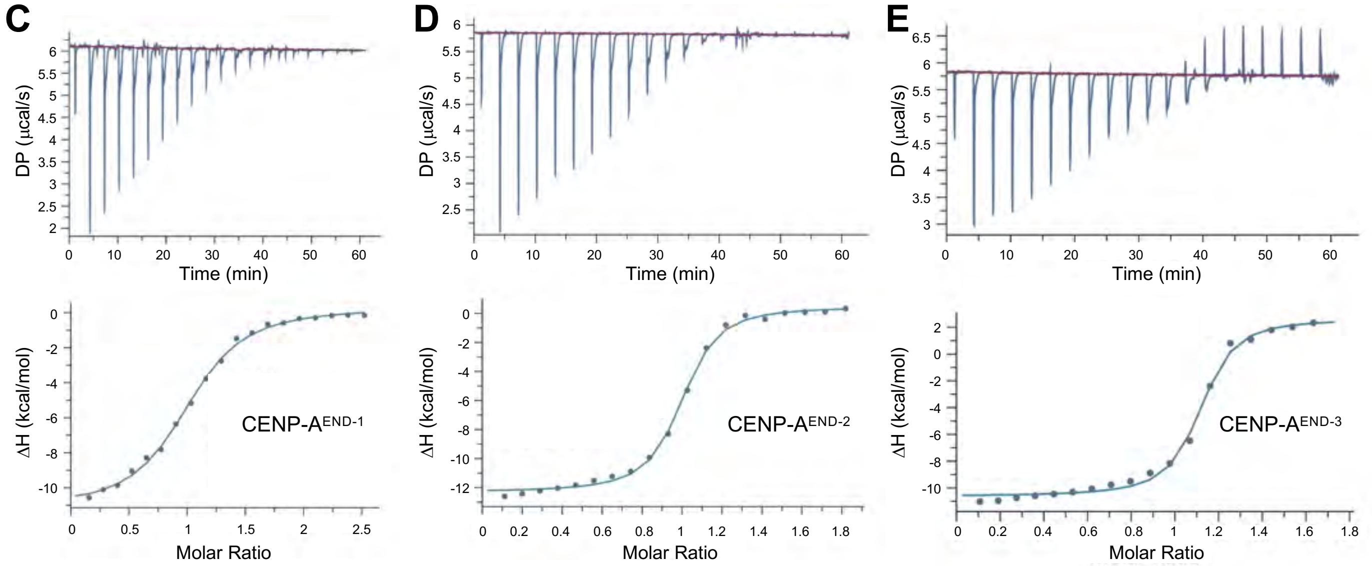

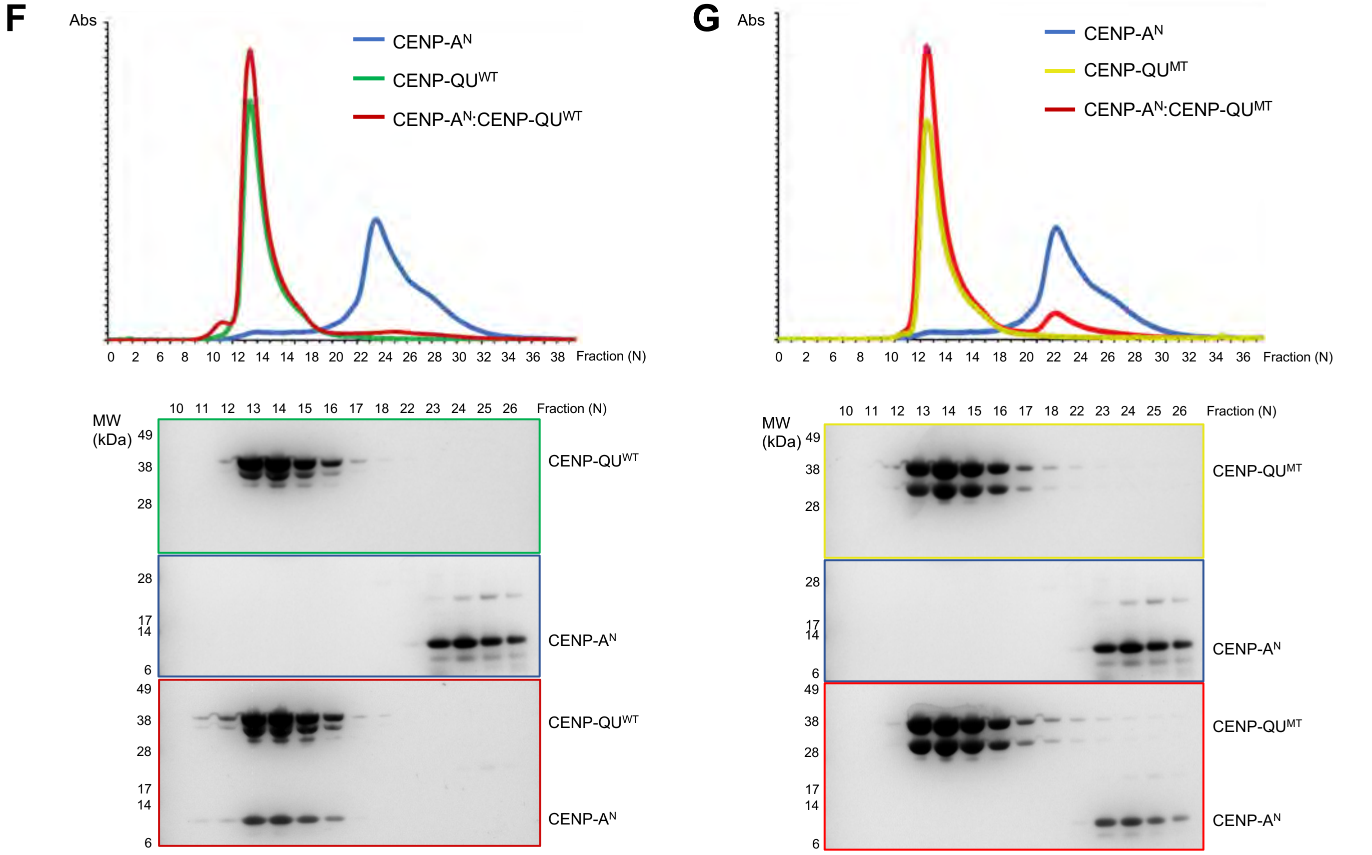

### Figure S8

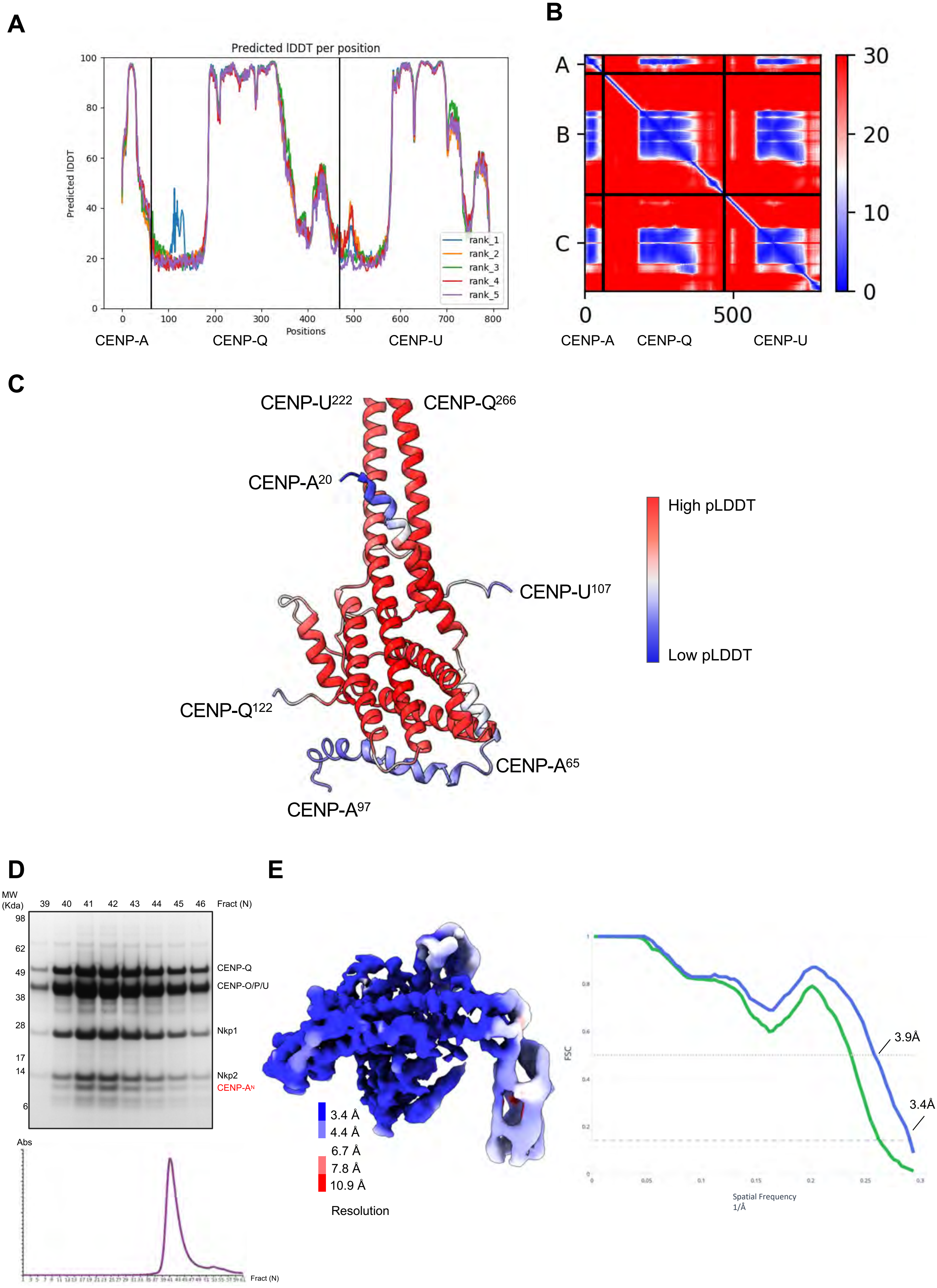

A
Predicted lDDT per position
Predicted lDDT
Positions
rank_1
rank_2
rank_3
rank_4
rank_5
CENP-A
CENP-Q
CENP-U
B
A
B
C
0
500
30
20
10
0
CENP-A
CENP-Q
CENP-U
C
CENP-U222
CENP-Q266
CENP-A20
CENP-U107
CENP-Q122
CENP-A65
CENP-A97
High pLDDT
Low pLDDT
D
MW (Kda)
39 40 41 42 43 44 45 46
Fract (N)
98
62
49
38
28
17
14
6
CENP-Q
CENP-O/P/U
Nkp1
Nkp2
CENP-AN
Abs
Fract (N)
E
3.4 Å
4.4 Å
6.7 Å
7.8 Å
10.9 Å
Resolution
FSC
3.9Å
3.4Å
Spatial Frequency
1/Å

### Figure S9

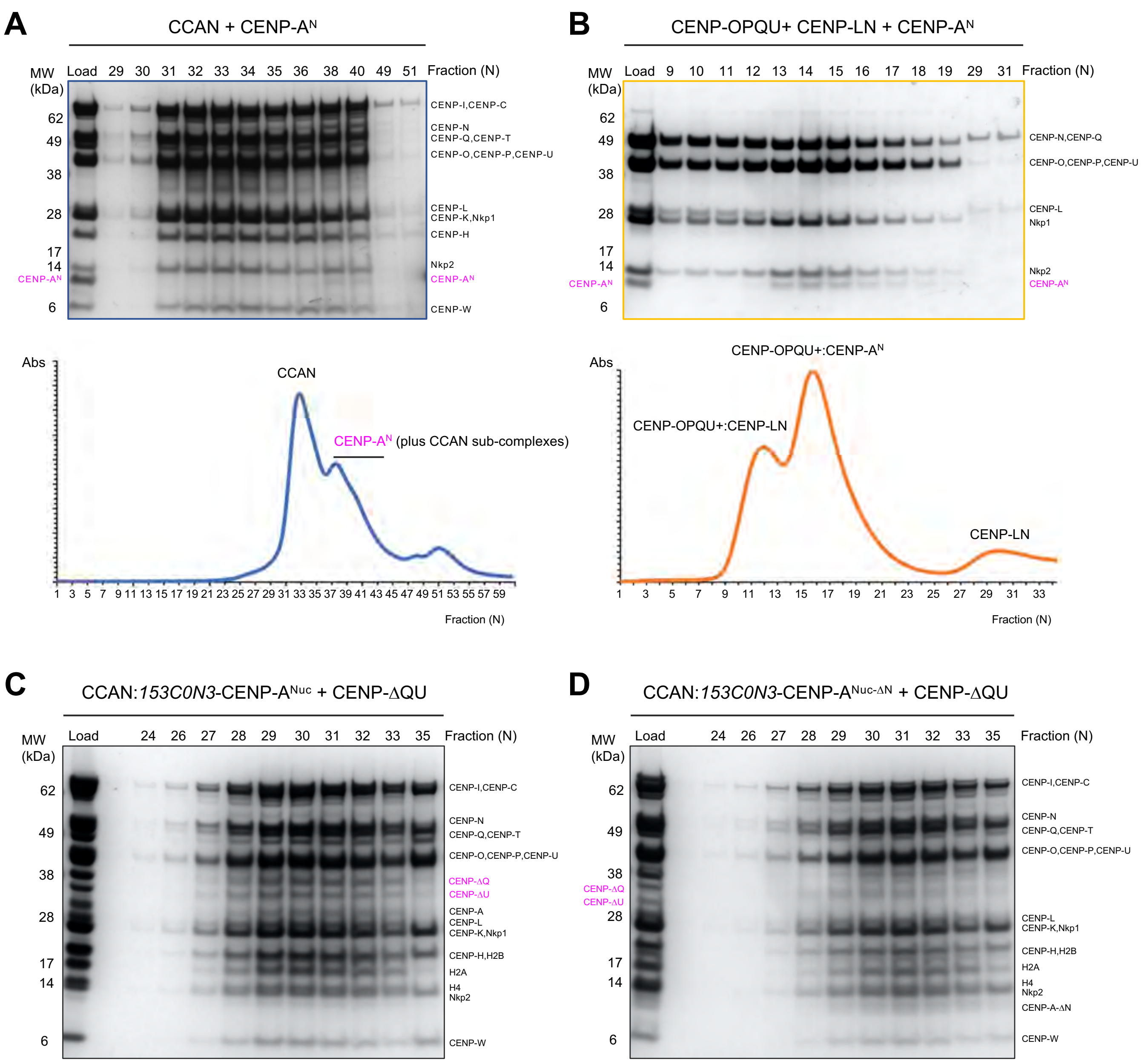

A
CCAN + CENP-A^N
MW (kDa)
Load 29 30 31 32 33 34 35 36 38 40 49 51 Fraction (N)
62
49
38
28
17
14
CENP-A^N
6
CENP-I,CENP-C
CENP-N
CENP-Q,CENP-T
CENP-O,CENP-P,CENP-U
CENP-L
CENP-K,Nkp1
CENP-H
Nkp2
CENP-A^N
CENP-W
Abs
CCAN
CENP-A^N (plus CCAN sub-complexes)
1 3 5 7 9 11 13 15 17 19 21 23 25 27 29 31 33 35 37 39 41 43 45 47 49 51 53 55 57 59
Fraction (N)
B
CENP-OPQU+ CENP-LN + CENP-A^N
MW (kDa)
Load 9 10 11 12 13 14 15 16 17 18 19 29 31 Fraction (N)
62
49
38
28
17
14
CENP-A^N
6
CENP-N,CENP-Q
CENP-O,CENP-P,CENP-U
CENP-L
Nkp1
Nkp2
CENP-A^N
Abs
CENP-OPQU+:CENP-A^N
CENP-OPQU+:CENP-LN
CENP-LN
1 3 5 7 9 11 13 15 17 19 21 23 25 27 29 31 33
Fraction (N)
C
CCAN:153C0N3-CENP-A^Nuc + CENP-ΔQU
MW (kDa)
Load 24 26 27 28 29 30 31 32 33 35 Fraction (N)
62
49
38
28
17
14
6
CENP-I,CENP-C
CENP-N
CENP-Q,CENP-T
CENP-O,CENP-P,CENP-U
CENP-ΔQ
CENP-ΔU
CENP-A
CENP-L
CENP-K,Nkp1
CENP-H,H2B
H2A
H4
Nkp2
CENP-W
D
CCAN:153C0N3-CENP-A^Nuc-ΔN + CENP-ΔQU
MW (kDa)
Load 24 26 27 28 29 30 31 32 33 35 Fraction (N)
62
49
38
CENP-ΔQ
CENP-ΔU
28
17
14
6
CENP-I,CENP-C
CENP-N
CENP-Q,CENP-T
CENP-O,CENP-P,CENP-U
CENP-L
CENP-K,Nkp1
CENP-H,H2B
H2A
H4
Nkp2
CENP-A-ΔN
CENP-W
